# Neural dynamics of shifting attention between perception and working-memory contents

**DOI:** 10.1101/2024.02.14.580280

**Authors:** Daniela Gresch, Sage E.P. Boettcher, Chetan Gohil, Freek van Ede, Anna C. Nobre

## Abstract

In everyday tasks, our focus of attention shifts seamlessly between contents in the sensory environment and internal memory representations. Yet, research has mainly considered external and internal attention in isolation. We used magnetoencephalography to compare the neural dynamics of shifting attention to visual contents within vs. between the external and internal domains. Participants performed a combined perception and working-memory task in which two sequential cues guided attention to upcoming (external) or memorised (internal) sensory information. Critically, the second cue could redirect attention to visual content within the same or alternative domain as the first cue. Multivariate decoding unveiled distinct patterns of human brain activity when shifting attention within vs. between domains. Brain activity distinguishing within- from between-domain shifts was broadly distributed and highly dynamic. Intriguingly, crossing domains did not invoke an additional stage prior to shifting. Alpha lateralisation, a canonical marker of shifting spatial attention, showed no delay when cues redirected attention to the same vs. alternative domain. Instead, evidence suggested that neural states associated with a given domain linger and influence subsequent shifts of attention within vs. between domains. Our findings provide the first insights into the neural dynamics that govern attentional shifts between perception and working memory.

**Significance Statement:** During almost every natural behaviour, our attention regularly shifts between sensory and memory contents. Although the systems and mechanisms of attentional control and modulation within the external and internal domains have been heavily studied in isolation, how attention crosses between these domains remains uncharted territory. Here, we provide the first study to investigate brain dynamics associated with shifting attention between contents in the sensory environment and memory representations. Using a novel experimental design, we isolated the patterns and dynamics of brain activity associated with shifting attention within vs. between the external and internal domains. Our findings reveal early, dynamic, and distributed patterns of activity that distinguish within- from between-domain shifts, offering fascinating initial insights, and opening new questions for investigation.

## Introduction

Attention can be selectively directed towards sensory information within the external environment or internal representations in memory, phenomena referred to as external and internal attention, respectively (1). During almost every natural behaviour, our attention fluctuates seamlessly between these attentional domains, rapidly transitioning focus between sensory and mnemonic contents. Despite playing an integral role in daily cognition, the mechanisms underlying attentional shifts between perceptual and memory contents have been relatively overlooked.

A large body of research has focused on studying isolated shifts of attention within perception (2, 3) or within working memory (4–6). Some investigations have also considered how shifts within each domain compare, emphasising shared neural systems and dynamics as well as highlighting dissociations (7–15). These studies have typically employed informative pre-cues or retro-cues in their task design to isolate brain activity specifically associated with shifting attention in either domain (4, 8–10, 14, 15).

In contrast, the investigation of between-domain shifts – transitioning attention between perceptual and mnemonic contents – is still at a nascent stage. A limited number of behavioural studies have highlighted higher costs when shifting across different attentional domains compared to shifting within the same domain (16–21). Brain-imaging and electrophysiological studies comparing task phases that rely more heavily on sensory stimuli vs. internal states have indicated that the lateral prefrontal cortex contributes to shifting attention between the external and internal domains (22–24). Complementary lines of research have suggested the hippocampus (25, 26), fronto-insular cortex (27), or retro-splenial cortex (28) may serve as interfaces between perception and memory. However, it is important to note that the task designs of these studies did not specifically contrast attentional shifts within and between domains. Beyond these initial clues, the dynamic neural mechanisms of cross-domain attention shifts are all to be discovered.

We aimed to expand upon these initial steps by investigating the brain dynamics associated with shifting attention between the perceptual and working-memory domains and by directly comparing these to the dynamics linked to shifting attention within domains. Building on our recent behavioural study (17), we used a combined perception and working-memory task in which informative cues indicated the location of sensory or mnemonic contents for subsequent report. Successive cues necessitated shifts of attention either within or between perception and working memory. We recorded magnetoencephalography (MEG) to track brain activity with high temporal resolution while participants performed the task and used multivariate pattern analysis to compare activity related to attentional shifts of different types. Based on our and other previous findings of performance costs for shifting attention between relative to within domains (16–21), we hypothesised the engagement of distinct processes when shifting attention between domains.

To preview our results, we demonstrate that dissociable patterns of brain activity accompany shifts between perception and working memory as opposed to shifts within either of these domains. These unique patterns of brain activity are broadly distributed, temporally dynamic, and occur without interfering with the timing of spatial attention shifts as indexed by the lateralisation of posterior 8-12-Hz alpha activity. In addition, we show distinct patterns of brain activity are associated with external and internal attention states themselves. These findings provide important initial clues about the dynamic neural bases that govern shifts of attention between perception and working-memory contents.

## Results

Twenty-five healthy human volunteers performed a combined perception and working-memory task (Fig. 1a). Two separate bilateral arrays each containing two randomly oriented bars were presented, with one array occurring before and the other after one or two spatially informative colour cues. That is, two bars appeared in an early, internal working-memory array, encoded before the cues, and two bars in a later, external perceptual array, appearing after the cues. At the end of each trial, participants reproduced the orientation of the bar that was last cued. Cues indicating the location of a previously encoded bar guided attention internally (i.e., retro-cues, internal cues), instructing which item in working memory was relevant for reporting. In contrast, cues indicating a previously unoccupied location directed attention externally (i.e., pre-cues, external cues), instructing which item in the upcoming perceptual array would be relevant. Half of the trials were no-shift trials in which only a single cue was presented, and participants reported the initially cued item. These randomly intermixed no-shift trials were included only to ensure that participants relied on the initial cue when performing the task. In the remaining trials, participants were tasked with shifting their attention to another item, which could be either an external or internal item, as indicated by a second cue. These double-cue trials were of central interest. The second cue always pointed to another item in the opposite hemifield but, importantly, equally often indicated that attention needed to be reoriented to an item of the same domain (i.e., within-domain shift) or an item in the other domain (i.e., between-domain shift). This resulted in four possible shift conditions: external-to-external, internal-to-internal, external-to-internal, and internal-to-external. The orthogonal manipulation of the to-be-reported target domain (external vs. internal) and the shift type (within-domain shift vs. between-domain shift) enabled us to investigate these two factors independently. The first cue in single-cue trials and the second cue in double-cue trials were 100% instructive and, thus, always indicated the target for report.

**Fig. 1.**
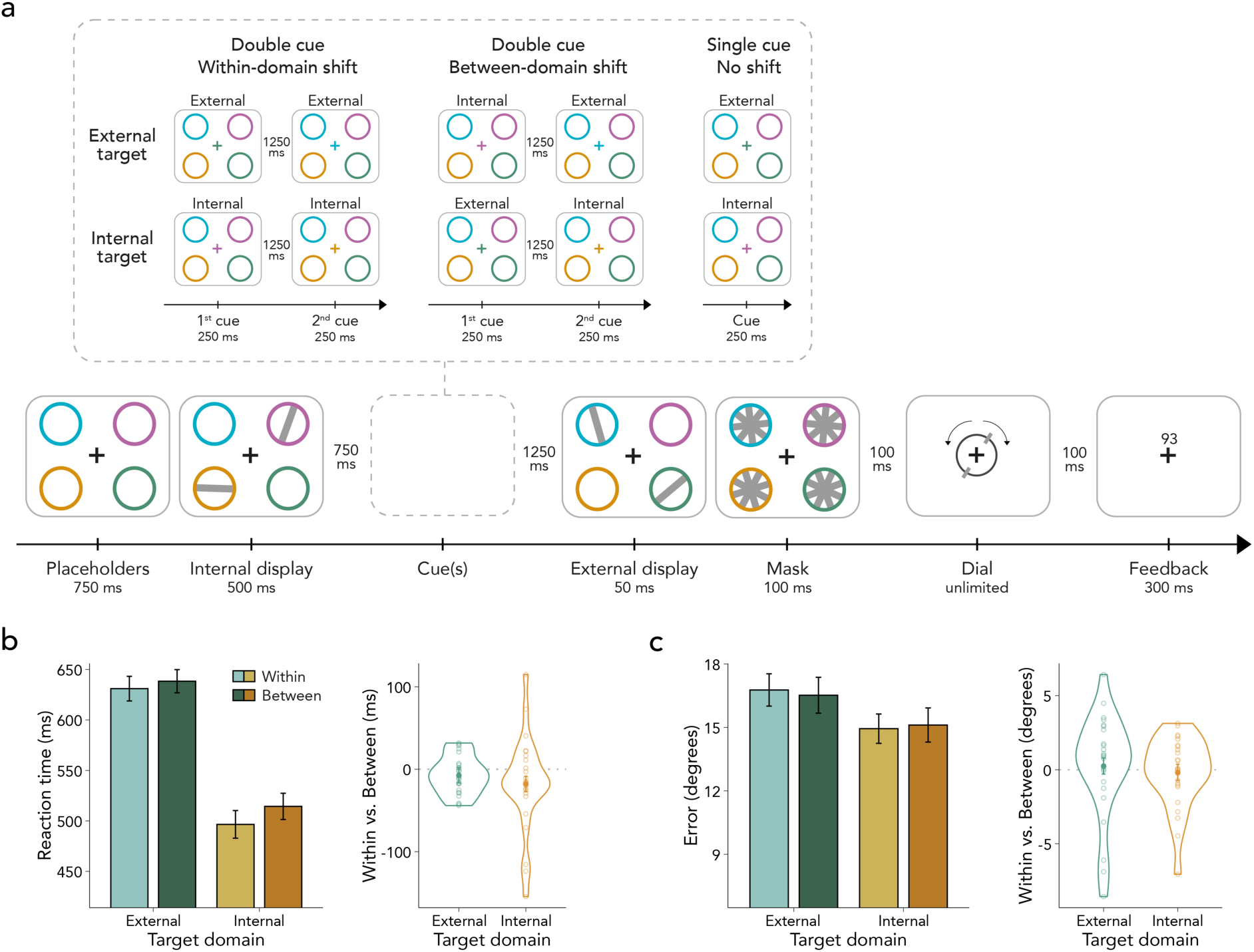
Task schematic and behavioural results. **(a)** Tilted bars were presented across a working-memory array, encoded before the cues (i.e., internal display), and a perceptual array, appearing after the cues (i.e., external display). In double-cue (i.e., shift) trials, two cues appeared between the presentation of the internal and external display. At the time of the second cue, attention needed to be reoriented either to an item of the same domain (i.e., within-domain shift) or to an item in the other domain (i.e., between-domain shift). To ensure participants relied on the first cue, 50% single-cue trials were included in which only one cue was presented, and thus no attentional reallocation was required. **(b)** Left panel: Average reaction times (RTs) as a function of target domain and shift type. RTs were faster for internal than external targets and for within- than between-domain shifts. The interaction between target domain × shift type was not significant. Error bars represent *SEM*. Right panel: Shift cost in RTs defined as the difference between shift types (i.e., within minus between) for each target domain. Dots represent individual participants. (c) Left panel: same as left panel in (b), but for reproduction errors. There were neither significant main nor interaction effects. Right panel: same as right panel in (b), but for reproduction errors.

### Shifting Attention Between Domains Incurs a Behavioural Cost

Our analysis focused on double-cue trials, as they uniquely enabled us to compare within- and between-domain shifts. Figure 1b depicts the average response-initiation times (RTs) relative to dial onset as a function of the target domain and shift type. We found longer RTs when reorienting attention between domains compared to reorienting within the same domain (*F*_(1,24)_ = 6.466, *p* = 0.018, η^2^_G_ = 0.003). This between-domain shift cost replicates previous behavioural findings (16–21). Moreover, RTs to the internal targets were faster than to external targets (*F*_(1,24)_ = 44.240, *p* < 0.001, η^2^_G_ = 0.216), likely because behavioural responses to the former could be prepared ahead of the response stage (29, 30). The interaction between target domain and shift type was not significant (*F*_(1,24)_ = 1.280, *p* = 0.269, η^2^_G_ < 0.001), suggesting that the effect of shift type did not depend on target domain (i.e., whether participants reported an internal or an external item).

As shown in Figure 1c, reproduction errors were similar in magnitude, yielding no significant main effect (target domain: *F*_(1,24)_ = 0.729, *p* = 0.402, η^2^_G_ = 0.008; shift type: *F*_(1,24)_ = 0.307, *p* = 0.585, η^2^_G_ < 0.001) or interaction (*F*_(1,24)_ = 1.552, *p* = 0.225, η^2^_G_ = 0.002).

### Distinct Patterns of Brain Activity for Within- vs. Between-domain Shifts

Our main aim was to investigate whether and when within- and between-domain shifts evoked reliably different neural activity patterns. To this end, we applied a time-resolved cross-validation decoding approach to the broadband sensor-level MEG data collected while participants performed the task. We first trained a Linear Discriminant Analysis (LDA) classifier to distinguish external-to-external and internal-to-internal trials (i.e., within-domain shifts) from external-to-internal and internal-to-external trials (i.e., between-domain shifts) on a timepoint-by-timepoint basis. We used the Area Under the Curve of the Receiver Operating Characteristic Curve (ROC-AUC) as the evaluation metric (31). The orthogonal manipulation of the target domain, shift type, and cued location ensured that the decoding of within- vs. between-domain shifts could be isolated, without being influenced by merely detecting the currently cued attentional domain or cued location.

We found reliable decoding of within- vs. between-domain shifts. As expected, no decoding of reorienting attention within vs. between domains was observed until after the second cue (Fig. 2a). Within- vs. between-shift decoding reached significance approximately 360 ms after the second-cue onset and persisted until external display presentation (∼1860-3000 ms, cluster *p* < 0.001). Complementary univariate analyses of event-related fields (ERFs) were insufficient to detect differences between shift types (Appendix SI, Fig. S1). This suggests that the reported multivariate differences may be driven by the pattern of activity rather than the amplitude of evoked responses.

**Fig. 2.**
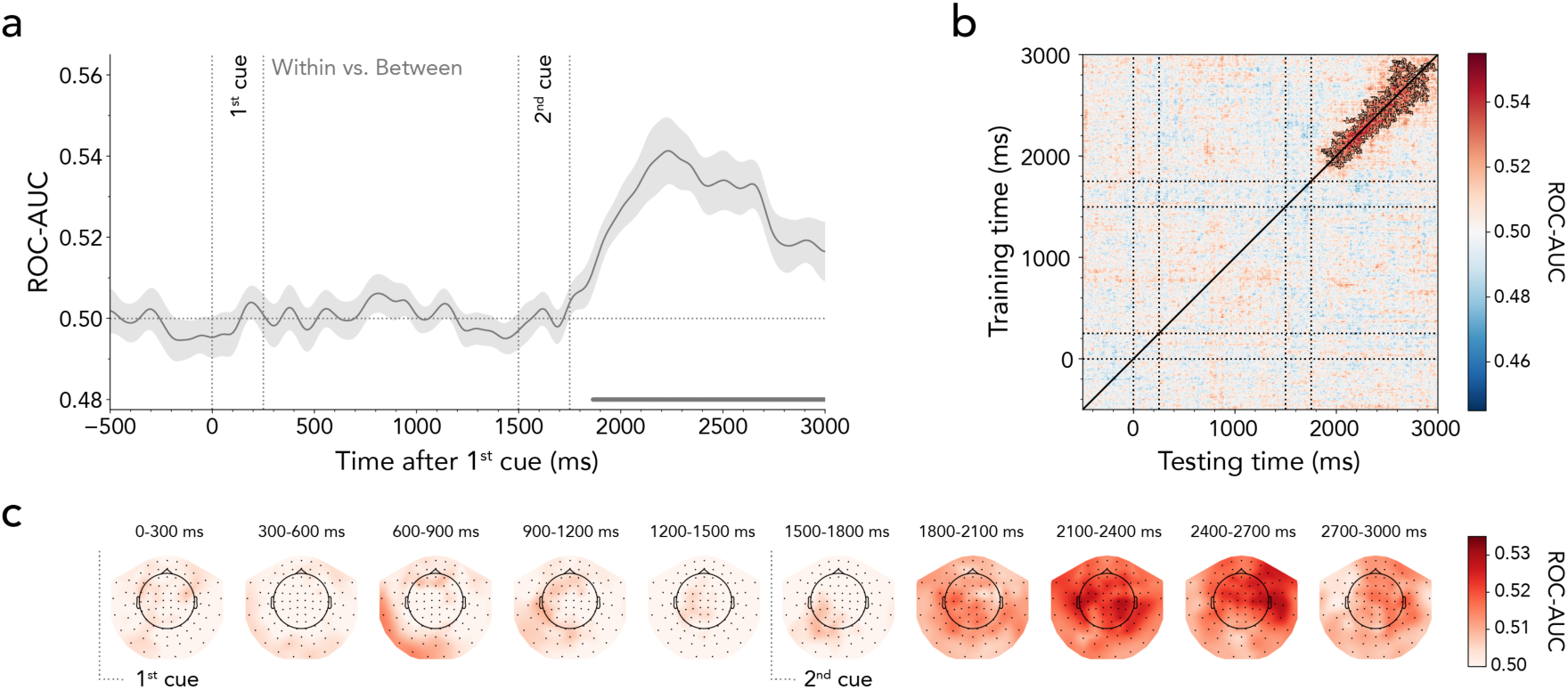
Time-resolved MEG decoding of within- vs. between-domain shifts of attention. **(a)** Average classifier performance for discriminating within- from between-domain shifts plotted over time. Time course shows *M* ± *SEM* across participants. Cluster-permutation corrected significant time points are indicated with horizontal lines. **(b)** Temporal generalisation matrix reflecting classifier performance in discriminating the two shift types. The diagonal corresponds to the time course illustrated in (a). Black outlines indicate significant decoding-clusters. **(c)** The MEG topography averages in 300-ms steps showing which sensors most strongly contribute to the overall decoding performance.

Since involuntary eye movements may contaminate MEG recordings and lead to decodable non-brain signals (32, 33), we trained a classifier to distinguish within- vs. between-domain shifts, but this time using only the x- and y-coordinates of the eye-tracking data. We could not decode the shift type based on the eye-tracking signal (Appendix SI, Fig. S2a). Moreover, we assessed whether there was mutual information between the eye-tracking and the MEG data. Appendix SI, Figure S2b shows the mutual information, with values of zero indicating that the MEG and eye-position signals are independent and larger values indicating a greater relationship between them. There was almost no mutual information between the eye-tracking and the MEG data when decoding within- vs. between-domain shifts. However, two participants were outliers, exhibiting stronger mutual information than the rest (Appendix SI, Fig. S2c). We therefore repeated the MEG decoding analysis while excluding these participants. This revealed that MEG-based decoding was not driven by participants with strong mutual information (Appendix SI, Fig. S2d).

To assess if the pattern of neural activity underlying the classification performance was stable or evolved dynamically over time, we performed a generalisation-across-time analysis (34, 35). Significant points off the diagonal would indicate that the classifier, when trained on data from time point A, can generalise to data from time point B. In our case, the evolving pattern of neural activity that distinguished within- from between-domain attention shifts was highly dynamic. The classifier only significantly generalised over brief periods (Fig. 2b; 1^st^ cluster *p* < 0.001, 2^nd^ cluster *p* = 0.027). This indicates that although we can consistently distinguish between trials in which participants shifted attention within vs. between domains, the neural code driving this distinction varied moment-to-moment. Our primary decoding analysis included data from all MEG sensor locations as features in the classifier. A searchlight analysis (i.e., iterative decoding for subsets of MEG sensors) was used to characterise the scalp distribution of sensors sensitive to within vs. between domain shifts. Figure 2c shows that the neural patterns distinguishing within- vs. between-domain shifts were broadly distributed.

In an exploratory analysis, we tested whether patterns of brain activity discriminating within- from between-domain shifts impacted later behavioural performance. To this end, we obtained single-trial decoding scores for each participant within the 360 to 1500 ms interval following the second cue (i.e., the significant cluster interval shown in Fig. 2a) and categorised these trial scores into low vs. high decoding performance. Appendix SI, Figure S3 depicts RTs as a function of shift type (within vs. between) and median split (low shift-type decoding vs. high shift-type decoding). Pairwise comparisons revealed a promising trend, showing a significant between-domain shift cost only when comparing trials with high classifier performance (*t*_(24)_ = 2.691, *p* = 0.013, *d* = 0.538). In contrast, there was no significant difference between shift types when considering trials with low classifier performance (*t*_(24)_ = 0.243, *p* = 0.810, *d* = 0.049). However, the direct comparison of the shift cost between low and high shift-type decoding did not produce significant results (*t*_(24)_ = 1.824, *p* = 0.081, *d* = 0.365). Thus, while the results tentatively suggest that the ability to decode within- vs. between-domain shifts in attention may impact later behavioural performance, these findings are not yet conclusive.

Together, our data show distinct patterns of brain activity associated with within- vs. between-domain shifts and reveal that these patterns occur early, are highly dynamic, are broadly distributed, and may impact later behaviour.

### Decoding External vs. Internal States Associated with the First and Second Cues

Having uncovered evidence for distinct patterns of brain activity during shifts of attention within vs. between domains, we investigated whether and when the brain differentiated the attended domain before and after the shifts occurred. As we noted previously, owing to our experimental design, decoding of the currently attended domain (external vs. internal) was orthogonal to the above decoding of within- vs. between-domain shifts.

We started our investigation by decoding external vs. internal attentional states, based on the labels of the first cue. To this end, we trained a classifier to distinguish external first-cue trials (i.e., external-to-external, external-to-internal) from internal first-cue trials (i.e., internal-to-internal, internal-to-external). Figure 3a shows successful decoding of the attentional domain from approximately 310 ms after first-cue onset (cluster *p* < 0.001). Interestingly, the significant decoding period extended into the second-cue phase, revealing that information about the initially cued attentional state persisted throughout the delay following the second cue.

**Fig. 3.**
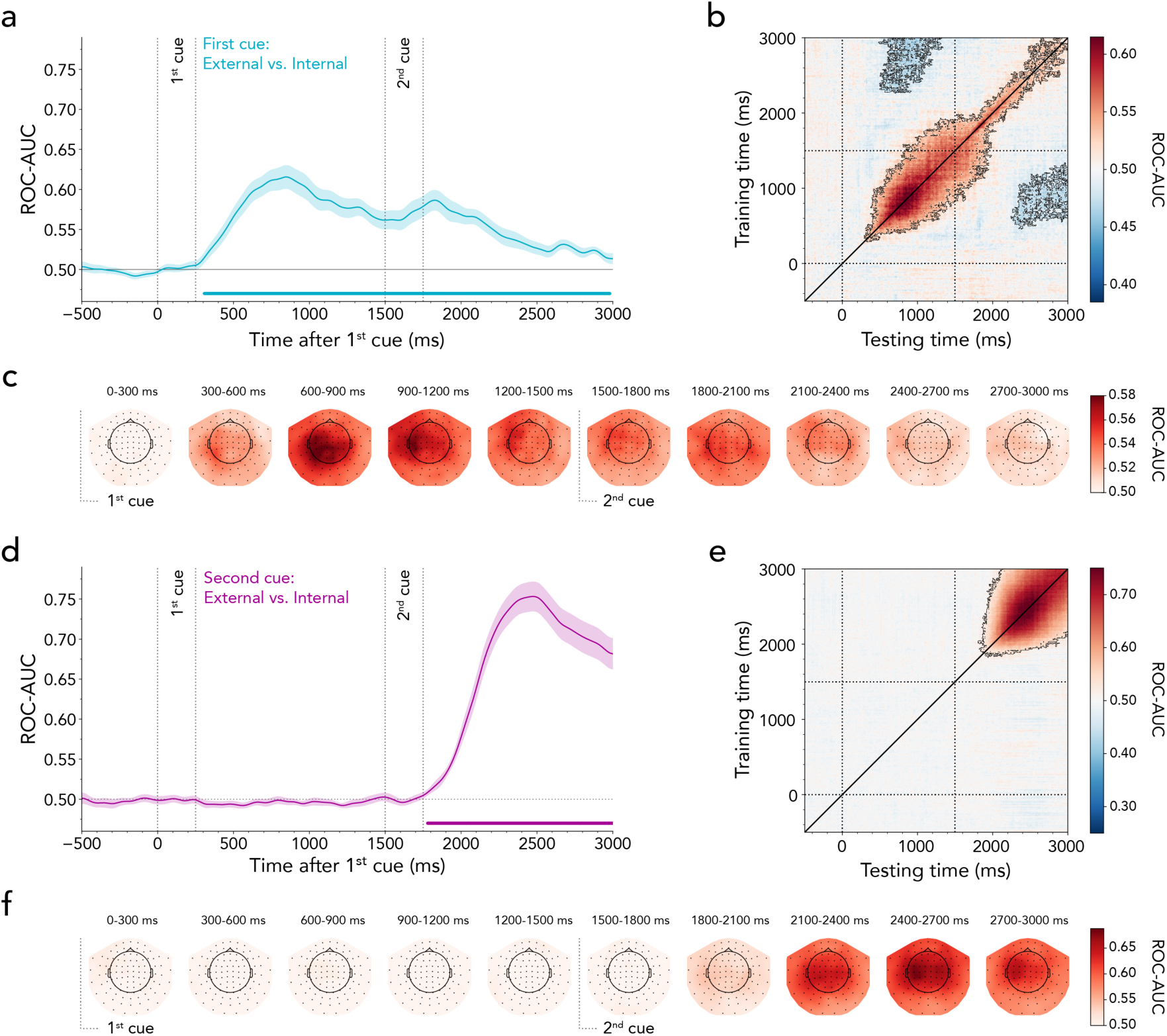
Time-resolved MEG decoding of external vs. internal attention. **(a)** Average classifier performance for distinguishing external from internal first cues plotted over time. Time course shows *M* ± *SEM* across participants. Cluster-permutation corrected significant time points are indicated with horizontal lines. **(b)** Temporal generalisation matrix reflecting classifier performance in discriminating the first-cued attentional domain. The diagonal corresponds to (a). Black outline indicates significant decoding clusters. **(c)** The MEG topography averages in 300-ms steps showing which sensors most strongly contribute to the overall decoding performance in discriminating external- and internal first cues. **(d)** Same as (a) but based on second-cue labels. **(e)** Same as (b) but based on second-cue labels. **(f)** Same as (c) but based on second-cue labels.

Temporal-generalisation analysis unveiled an off-diagonal shape of generalisation following the first cue, indicating temporal stability of the neural pattern evoked by the initially cued domain (Fig. 3b; cluster *p* < 0.001). In contrast, after the second cue, decoding performance occurred mainly along the diagonal, with a narrower spread to neighbouring timepoints. In addition, we found off-diagonal below-chance decoding (i.e., the classifier systematically guessed incorrectly; 1^st^ cluster *p* = 0.024, 2^nd^ cluster *p* = 0.025). The searchlight decoding analysis demonstrated that the contribution of sensors to the external vs. internal decoding was broadly distributed, with peaks observed in the posterior and left-central sensors occurring between 600 and 1200 ms (Fig. 3c).

To determine whether the decoding of the first-cued attentional domain was predominantly influenced by a particular trial type, we employed two additional classifiers: one to distinguish between external-to-external and internal-to-external shifts, and another to distinguish between internal-to-internal and external-to-internal shifts. Appendix SI, Figure S4a shows that classifier performance did not differ between these two decoding analyses, thus resembling the general external- vs. internal- domain decoding time course shown in Figure 3a. In addition, we again took precautions to ensure that eye movements did not drive the external vs. internal first-cue MEG decoding results. The strength of the decoding based on gaze position was relatively weak and far from sufficient to explain the effects when decoding brain activity (Appendix SI, Fig. S2e). Furthermore, exclusion of participants who exhibited the highest mutual information between eye position and MEG decoding did not alter the MEG decoding results (Appendix SI, Fig. S2f-h).

Next, we aimed to decode the domain indicated by the second cue. To this end, we trained a classifier to distinguish between external second-cue trials (i.e., external-to-external, internal-to-external) vs. internal second-cue trials (i.e., internal-to-internal, external-to-internal). Our results revealed significant decoding of the second-cued attentional domain from approximately 280 ms after second-cue onset (i.e., 1780 ms after first-cue onset; cluster *p* < 0.001; Fig. 3d). As intended by our design, the second-cued domain could not be decoded before the onset of the second cue. The temporal-generalisation matrix in Figure 3e showed a temporal spread, implying some stability in the activation pattern (cluster *p* < 0.001). Similar to the first-cue analysis, the searchlight analysis indicated a distributed pattern of sensor contribution to decoding external vs. internal attentional domains (Fig. 3f).

To ensure that the results of the domain-decoding were not driven by the contrast between specific trial types, we trained one classifier to discriminate between external-to-external and external-to-internal shifts and another one to discriminate between internal-to-internal and internal-to-external shifts. While both decoding time courses reached significance following the second cue, the internal-to-internal vs. internal-to-external contrast reached significance earlier than the external-to-external and external-to-internal contrast (Appendix SI, Fig. S4b). These findings revealed earlier domain differentiation when initially cued about the internal domain compared to when previously cued about an external item. In line with the first-cue analysis, we ensured that the second-cue MEG decoding results were not dependent on eye movements (Appendix SI, Fig. S2i-l).

### Lingering First-cued Attentional Domain Is Linked to Within- vs. Between-domain Shifting

Our results demonstrate that attentional domains indicated by the first and second cue could be decoded. Interestingly, the significant decoding of the first-cued attentional state persisted even after the second domain was cued. Supplementary analyses revealed that during the period following the second cue, the decodability of the continued information concerning the first cue was not contingent on the quality of processing of the second cue (Appendix SI, Fig. S5). Hence, this finding suggests that lingering processing related to the attentional domain of the first cue and instantiating the attentional domain associated with the second cue proceeded in tandem. In other words, information regarding the past and current attentional state co-existed after the second cue.

Previous task-switching research has argued that switch costs can arise from interference from prior, but now task-irrelevant, information (36, 37). We, therefore, tested whether a similar interaction might arise between lingering activity related to the first-cued domain and the ability to decode within- vs. between-domain shifts. To this end, we median-split our data into trials with good vs. bad first-cued domain decoding following the onset of the second cue (i.e., 1500-3000 ms). Next, we trained a classifier to distinguish within vs. between-domain shifts in the two resulting data subsets. The within- vs. between-domain shift type decoding only yielded a significant cluster in trials with high decodability of the lingering first-cued domain (Fig. 4a; ∼1910-2890 ms, cluster *p* < 0.001). This was corroborated by a direct comparison, showing that trials with stronger lingering first-cue decodability had significantly more successful within- vs. between-domain decoding (∼2330-2870 ms, cluster *p* = 0.005). Figure 4b displays the within- vs. between-shift decoding topographies for low (top panel) and high (bottom panel) lingering first-cue decoding.

**Fig. 4.**
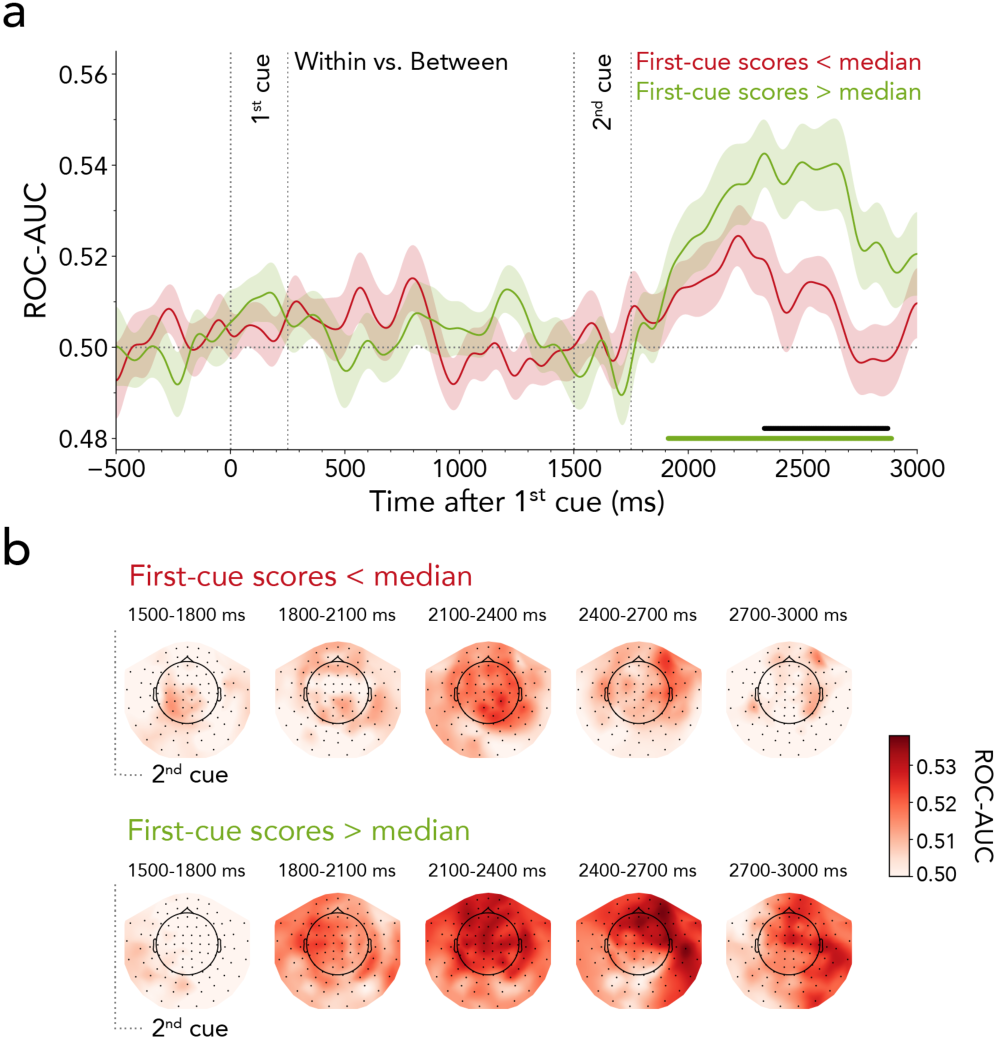
Relationship between lingering first-cued attentional domain and shift type. **(a)** Time-resolved average classifier performance for discriminating within- from between-domain shifts as a function of bad (red) vs. good (green) decoding of the lingering first-cued attention state. Time courses show *M* ± *SEM* across participants. Horizontal lines indicate significant clusters, black line denotes difference between decoding time courses. **(b)** MEG topography averages in 300-ms steps showing which sensors most strongly contribute to the overall decoding performance in discriminating within- and between-domain shifts. Top panel: trials with low decodability of the lingering first-cued domain. Bottom panel: trials with high decodability of the lingering first-cued domain.

In addition, we found tentative evidence suggesting that the decoding quality of the lingering first-cued domain influenced the behavioural between-domain shift cost (Appendix SI, Fig. S6). Our data revealed a significant shift cost only when comparing trials with high classifier performance (*t*_(24)_ = 2.494, *p* = 0.020, *d* = 0.499), while no difference emerged between shift types when considering trials with low classifier performance (*t*_(24)_ = 0.945, *p* = 0.354, *d* = 0.189). However, a direct comparison of the shift cost between low and high lingering-domain decoding did not yield significant results (*t*_(24)_ = 1.287, *p* = 0.210, *d* = 0.257).

Overall, our results suggest that the extent to which the first-cued state lingers is associated with the classifier’s ability to differentiate within- from between-domain shifts and may also predict subsequent behavioural shift costs.

### Spatial Shifts of External and Internal Attention as Tracked in Alpha Lateralisation

Thus far, we have investigated shifting from two essential perspectives. Firstly, we contrasted brain activity when transitioning between domains vs. within a single domain. Secondly, we examined whether brain activity distinguished between attentional domains following the first and second cues. Our task design additionally enabled us to investigate attentional shifts from a third perspective, involving the spatial orienting of attention between locations. We intentionally designed our task to guarantee that shifts occurred between items positioned in distinct hemifields, enabling us to investigate the lateralisation of MEG activity according to the location to which attention was shifted.

A canonical marker of spatial attention shifts is the lateralisation of posterior 8-12 Hz alpha activity. Contra- vs. ipsilateral attenuation of 8-12 Hz alpha oscillations has been linked to relative increases in excitability in sensory brain areas resulting from shifts of spatial attention in anticipation of relevant sensory stimuli as well as during selection of relevant working-memory content (14, 30, 38– 42). Therefore, we focused on alpha lateralisation to delineate spatial shifts of attention after external and internal first and second cues. These additional analyses complement our primary findings from the decoding analyses and help relate our work to the existing neuroscience literature on external and internal spatial shifts of attention.

Figure 5a and 5c show the time- and frequency-resolved lateralisation of spectral power in contra- vs. ipsilateral visual sensors for external and internal first cues, respectively. At the time of the second cue, participants were always required to shift attention from one hemifield to the other. Consequently, relative suppression of contralateral activity in response to the first cue was followed by relative enhancement of contralateral activity (i.e., a reversal of lateralisation) after second-cue onset (external first cued: 1^st^ cluster *p* < 0.001, 2^nd^ cluster *p* < 0.001; internal first cued: 1^st^ cluster *p* < 0.001, 2^nd^ cluster *p* = 0.004).

**Fig. 5.**
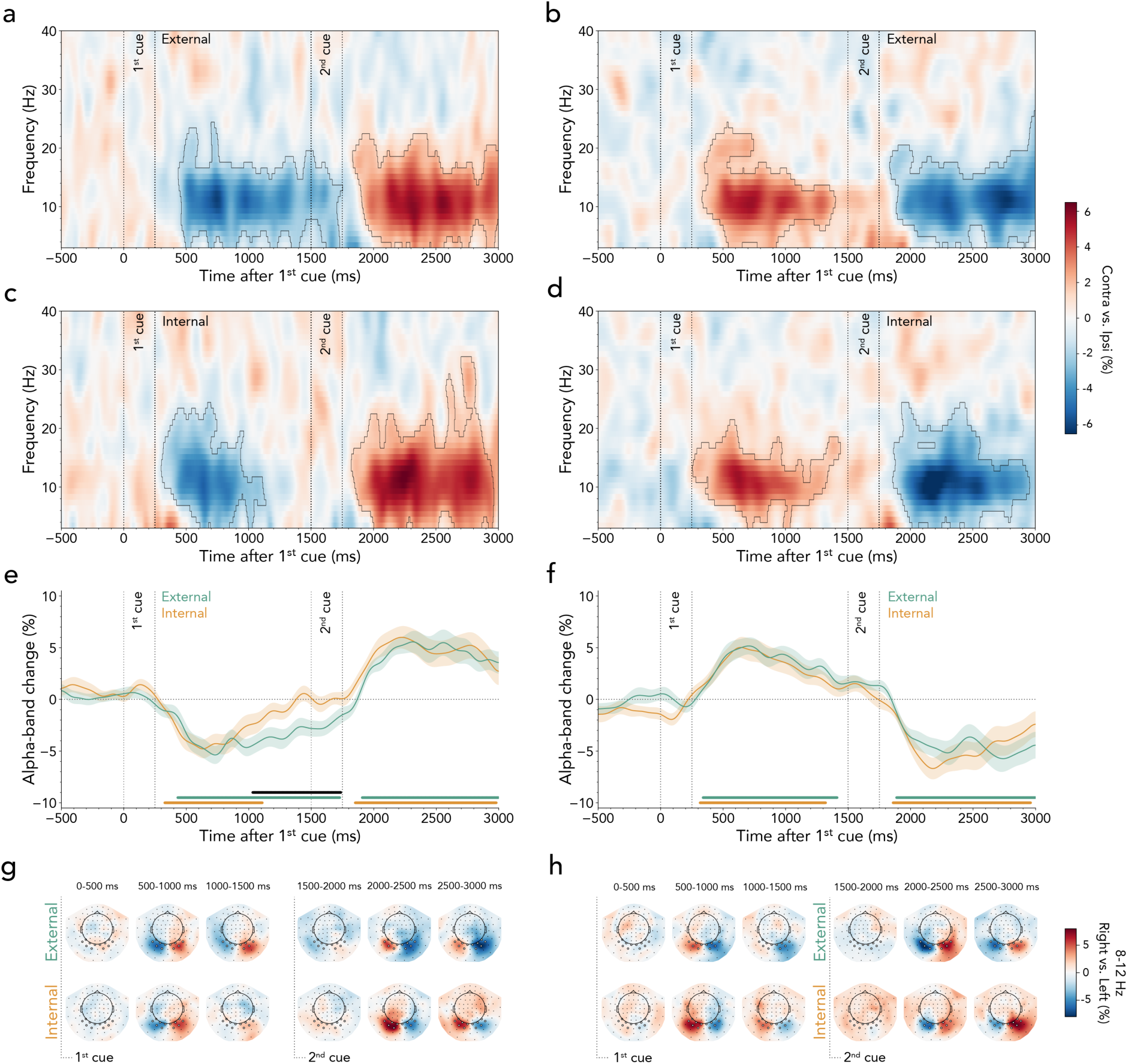
Lateralised neural activity following external and internal cues. **(a)** Time-frequency plots of neural lateralisation following external first cues. Plots show the difference in spectral power contra- vs. ipsilateral to the cued side, extracted over the predefined left and right sensor clusters. Black outlines indicate significant clusters. **(b)** Same as (a) but for external second cues. **(c)** Same as (a) but for internal first cues. **(d)** Same as (a) but for internal second cues. **(e)** Time courses of neural lateralisation in the 8-12 Hz alpha band relative to the first cue. Time courses show *M* ± *SEM* across participants. Horizontal lines indicate significant clusters, black line denotes difference between time courses. **(f)** Same as (e) but relative to second cues. There was no significant difference between time courses. **(g)** Topographies of power differences following right- vs. left-directing external first cues in the 8-12 Hz alpha band. Top panel shows topographies for external first cues, bottom panel for internal first cues. **(h)** Same as (g) but for second cues.

To chart the time course of alpha lateralisation, we averaged the time-frequency data along the classical alpha band (8-12 Hz; Fig. 5e). We confirmed the typical contra- vs. ipsilateral alpha attenuation following both external and internal first cues. Clusters of lateralised alpha attenuation alternated between hemispheres after the first (external first cued: ∼430-1730 ms, cluster *p* < 0.001; internal first cued: ∼330-1110 ms, cluster *p* < 0.001) and second cue (external first cued: ∼1910-3000 ms, cluster *p* < 0.001; internal first cued: ∼1850-2980 ms, cluster *p* = 0.004). The time course of the initial contra- vs. ipsilateral alpha attenuation was similar for both external- and internal first-cue trials. However, in line with prior research, the duration of the alpha modulation differed (14, 43). Lateralised alpha suppression was more transient following internally directing first cues and more sustained following externally directing first cues, as indicated by a significant cluster from approximately 1030 to 1740 ms after first-cue onset (cluster *p* < 0.001). A topographic inspection of external and internal first-cued trials confirmed a clear posterior topography for these effects (Fig. 5g, upper and lower panel, respectively).

The same time-frequency analysis was conducted based on the attentional domain associated with the second cue. Figure 5b and 5d illustrate that the relative suppression of contralateral activity in response to the second cue was preceded by a relative enhancement of contralateral activity (i.e., a reversed lateralisation) after first-cue onset (external second cued: 1^st^ cluster *p* < 0.001, 2^nd^ cluster *p* < 0.001; internal second cued: 1^st^ cluster *p* = 0.002, 2^nd^ cluster *p* = 0.002). As seen in the alpha-band time-courses (Fig. 5f), there was no difference between alpha-band lateralisation following the first cue (external second cued: ∼360-1400 ms, cluster *p* < 0.001; internal second cued: ∼360-1300 ms, cluster *p* = 0.002) or following the second cue (external second cued: ∼1920-3000 ms, cluster *p* = 0.003; internal second cued: ∼1920-2940 ms, cluster *p* < 0.001). Topographies showing the difference between trials with right- vs. left-cued items confirmed that these visual signatures were prominent in the corresponding right and left visual sensors (Fig. 5h, top panel: external cue, bottom panel: internal cue).

Together, these complementary analyses of alpha lateralisation reveal that attention lingers longer after the first cue when directed externally vs. internally, while re-orienting after the second cue proceeds similarly regardless of whether spatially re-orienting to an internal or external item.

### Alpha Lateralisation Proceeds Similarly During Within- and Between-domain Shifts

Given the effectiveness of lateralised alpha attenuation in tracking the spatial orienting and re-orienting of attention, we finally turn to the important question of whether spatial orienting following the second cue would be delayed when shifting attention between as compared to within domains. If shifting between domains engages additional processes prior to implementing a spatial shift, we would expect a delay in the onset of spatial re-orienting when shifting attention between domains.

Figure 6a depicts the time course of alpha lateralisation relative to the second cue, split by whether the second cue indicated a within- or a between-domain shift (within-domain shift: 1^st^ cluster ∼340-1300 ms, cluster *p* = 0.002; 2^nd^ cluster ∼1910-3000 ms, cluster *p* < 0.001; between-domain shift: 1^st^ cluster ∼370-1380 ms, cluster *p* < 0.001; 2^nd^ cluster ∼1860-3000 ms, cluster *p* < 0.001). Comparing time courses revealed no differences in lateralised alpha-band activity between shift types. Moreover, for both within- and between-domain shifts, source localisation confirmed that the difference between trials with right- vs. left-cued items originated in right and left posterior cortices (Fig. 6b, upper panel: within-domain shift, lower panel: between-domain shift). These findings suggest that the initiation of the spatial shift remained unaffected, despite our decoding analyses revealing distinguishable neural processing engaged during within vs. between-domain shifts.

**Fig. 6.**
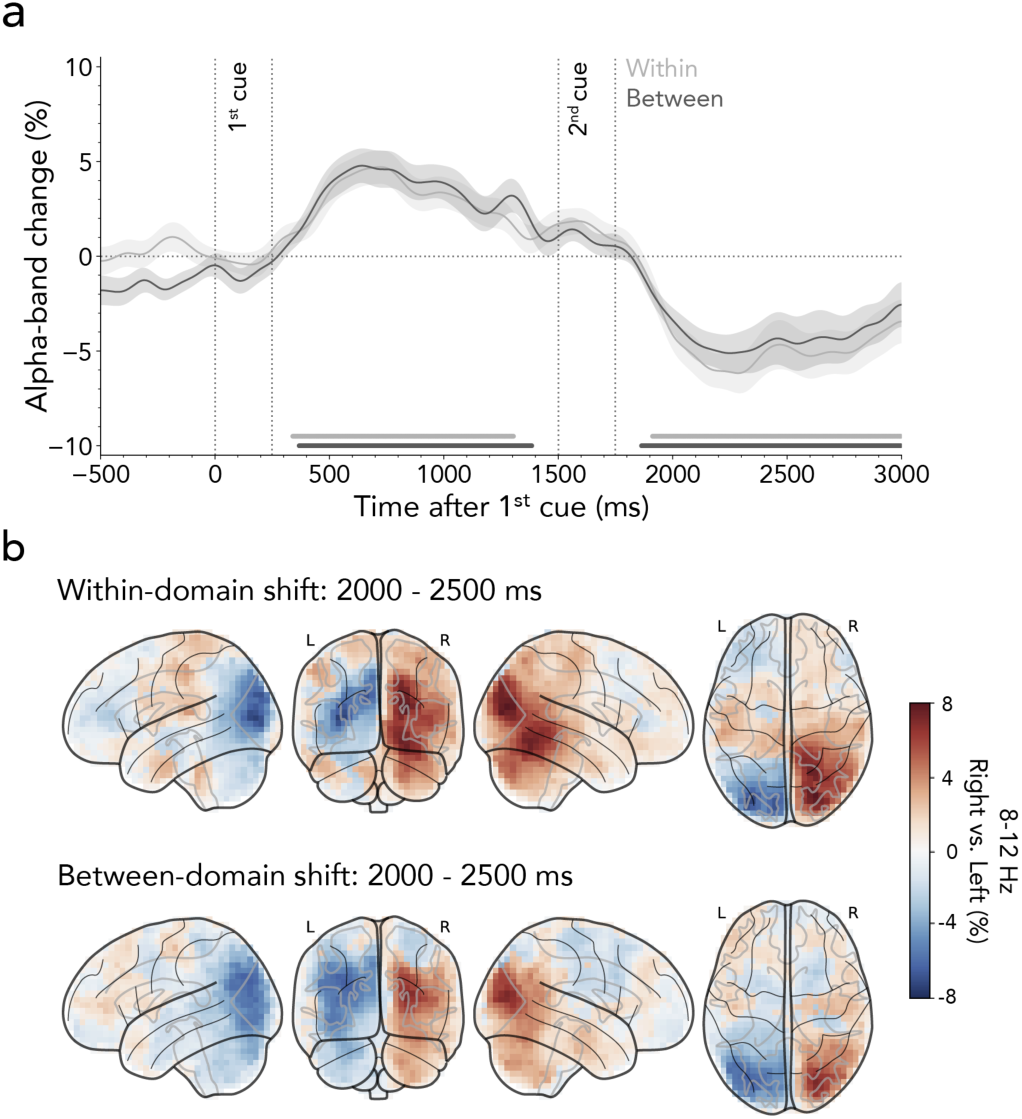
Lateralised neural activity during within- and between-domain shifts. **(a)** Time courses of neural lateralisation in the 8-12 Hz alpha band relative to second cues indicating within- vs. between-domain shifts. Time courses show *M* ± *SEM* across participants. Horizontal lines indicate significant clusters. **(b)** Participant-averaged voxelwise difference in alpha-band activity between trials in which cues indicated visual content in the right vs. left hemifield. The top panel shows source-reconstructed alpha-band activity during within-domain shifts, while the bottom panel illustrates activity during between-domain shifts.

This conclusion was further reinforced by the analysis of lateralised ERFs from posterior sensors, which revealed N2pc-like effects after both within- and between-domain shifts. These lateralised N2pc-like effects were equivalent for both shift types (Appendix SI, Fig. S7).

Taken together, our data provide no evidence for an additional stage invoked by between- domain shifts that must be resolved before attention can shift in space.

## Discussion

Our study uncovers direct neurophysiological evidence for distinct brain activity when shifting between perception and working-memory contents compared to when shifting between contents within either domain. The carefully balanced task design – which orthogonally manipulated shift type and target domain while also necessitating spatial shifts of attention – offered valuable insights into the dynamic neural bases that govern shifts between perception and working memory.

In principle, multiple scenarios can explain the behavioural costs and neural differences between shift types, and these need not be mutually exclusive. One obvious scenario is the addition of a supraordinate supervisory control process that acts upon the neural systems that control between-domain attentional shifts (22, 23, 44, 45). This supraordinate control mechanism could be invoked when initiating a shift between perception and working memory (and vice versa) but not when shifting between contents within either domain. Although it is most intuitive to consider a supraordinate control mechanism occurring when attention shifts between domains are initiated, it is also possible to imagine supraordinate control mechanisms operating at other stages of attention deployment. Attention comprises several functions to guide adaptive behaviour, such as disengaging, orienting, selecting, prioritising, and output gating (46, 47). Accordingly, additional supraordinate mechanisms could operate at any or multiple points along the way. An alternative type of scenario invokes no additional control process(es). Instead, the systems for controlling external vs. internal attention may coexist and compete for access to the relevant sensory and/or motor representations. In this scenario, the additional cost of shifting between perception and memory (and vice versa) would arise from resolving a stronger degree of competition between as compared to within domains. As we outline below, our findings offer important initial clues regarding the factors that distinguish within- and between-domain shifts.

We unequivocally demonstrated distinct neural processes during within- and between-domain shifts. By tracking neural dynamics, we showed that differences occur early following the second cue (Fig. 2), suggesting they are unlikely to emerge exclusively at the response stage. In line with this, our data provided tentative evidence suggesting that the degree to which shift types can be decoded may have behavioural relevance, impacting later between-domain shift costs in RTs (Appendix SI, Fig. S3). We cautiously argue, therefore, that previously observed between-domain shift costs in performance (16–20, 23, 48) are unlikely explained exclusively by interference at the late response-level stage, but rather emerge already during the attention-shifting process itself.

Interestingly, by analysing alpha-band lateralisation, we found no evidence indicating differences in visual activity when orienting spatial attention within vs. between domains (Fig. 6; see Appendix SI, Fig. S7 for lateralised ERFs). This observation challenges the intuitive scenario proposing that a supraordinate control mechanism is invoked when initiating an attentional shift that crosses domains. Such a scenario would predict a delayed shift of spatial attention. We found no evidence for this scenario. Spatial orienting and selection functions of attention proceeded with equivalent time courses, as reflected by lateralised modulations of posterior alpha and ERFs. These findings complement a similar observation that between-domain shifts did not alter patterns of gaze biases associated with spatial shifts (17), further suggesting that no additional, time-consuming gating step occurs before spatial shifts between domains.

Neural patterns differentiating within- vs. between-domain shifts evolved relatively dynamically and were broadly distributed (Fig. 2b). Differences between shift types are thus not solely attributable to a single tonic state that differs between shift types but likely result from the dynamic interplay of multiple states. Moreover, the distributed nature of the within- vs. between-shift decoding suggests that large-scale networks may be involved (Fig. 2c). It will be important for future brain-imaging investigations using functional Magnetic Resonance Imaging (fMRI) and/or animal models to substantiate this broad distribution and to pinpoint the precise spatial underpinnings of the here-reported findings.

It is useful to consider that each within- and between-domain shift is a consequence of the attended domains before and after the shift. Therefore, to understand the nature of the two shift types, it is also essential to investigate the neural dynamics evoked by the first- and second-cued attentional domains. While some studies have emphasised the shared neural systems and dynamics between selection mechanisms in external and internal attention (10, 15), others have highlighted important differences (7, 11, 46, 47, 49–53). Underscoring the differences, our data demonstrate successful decoding of external vs. internal attention (Fig. 3). Moreover, we show that decoding of the first-cued domain persisted even after the presentation of the second cue.

The carry-over decoding effect of the first-cued domain occurred regardless of the domain indicated by the second cue (Appendix SI, Fig. S4a). This finding highlights that the attentional state invoked by the second cue did not overrule or wipe out the lingering state invoked by the first cue. Multiple processes could have contributed to the lingering decoding of the first-cued domain, and we are unable to adjudicate among these without further investigation. For example, anticipating the spatial location or sensory features of a target in the external domain may have elicited sustained neural activity according to spatial or feature-based receptive field properties (54–56). In contrast, cueing internal representations in working memory often triggers more transient sensory modulation (14, 43). Our alpha-lateralisation findings hint at these effects, with more prolonged modulation following spatial cues indicating targets in the external than internal domains (Fig. 5a, c, e, g). Alternatively, the presentation of the second cue may have activated latent functional states in working memory through a pinging mechanism (57–59). This neural reactivation can also occur in the presence of active working-memory representations, causing an increase in their decodability (60). Although the second cue indicated a different item than the first cue, its associated sensory input may have interacted with latent or activated working-memory states, potentially acting as a ping in trials in which attention was initially directed to an internal item. Another possibility is that the first cue may have placed the system in different states of sensorimotor preparation. If an internal item was cued, its representation could have been readily transformed into a motor code (29, 30, 61), whereas this was not possible when sensory items were anticipated by external attention. Therefore, the sustained decoding of the first-cued attentional domain following the second cue could reflect a lasting effect of output gating following internal but not external first cues. Any, or all, of these differences may have contributed to the successful decoding of the first-cued domain after the second cue.

The decodability of the lingering first-cued attentional domain did not trade off against the decoding of the second-cued attentional domain (Appendix SI, Fig. S5). Instead, after the second cue, information processing unfolded for both the first- and second-cued attentional domains in tandem, implying the co-existence of preceding and current attention states in the brain. Though our data do not directly tap into the nature or potential functional relevance of these co-existing states, they did reveal that the degree to which the first state lingered (i.e., could still be decoded after the second cue) predicted how well our classifier could distinguish within- from between-domain shifts (Fig. 4). Moreover, our data offered tentative evidence that lingering activity from the first cue predicted the behavioural between-domain shift cost (Appendix SI, Fig. S6). This suggests that preceding attentional states influence upcoming attentional shifts. What role such lingering states play, and whether/how they mediate parallel processing of contents from preceding and current attention states remains an exciting endeavour for future studies.

Our findings and task design pave the way for investigating the dynamic mechanisms orchestrating the seamless shifts of attention between external and internal contents to guide adaptive flexible behaviour. We offer an initial exploration, opening the door to relevant unanswered questions and encouraging further task manipulations. Many additional and interesting factors and dimensions remain to be considered. Additional manipulations will yield complementary insights and contribute to a more comprehensive understanding of the principles and mechanisms that govern attention shifts between perception and working memory in service of adaptive behaviour.

One important factor to consider is that external and internal attention can be operationalised in different ways. In our study, we exploited pre- and retro-cues as tools to orient attention towards anticipated near-future external stimuli or near-past sensory stimuli maintained in working memory, respectively. In this conceptualisation and task design, participants can select relevant contents from working memory when cued internally but must await the incoming stimulation before selecting contents when cued externally. However, external attention is not limited to anticipating sensory events but also operates when sensory content is available for immediate selection, such as during visual search. In future studies, it will be interesting to investigate the dynamics of transitioning between external and internal content in tasks where external information is presented concurrently with external cues and remains perceptually available until the reporting stage.

Moreover, the informative cues used in this study engaged goal-directed processes. The source of attentional control was thus inherently endogenous (i.e., internal) for both externally and internally directed cues. The core distinction separating external from internal attention concerned the stimulus contents to be selected and prioritised: upcoming sensory stimulation or maintained memory representations. Future research can also compare how various sources of attention guide between-domain shifts.

Lastly, in our study, all double-cue trials required a spatial shift of attention when the second cue was presented. This design characteristic ensured that external and internal attention conditions were optimally balanced and minimised potential interference from items occurring at the same location. Future studies can investigate whether and how spatial overlap and proximity interact with shifting attention between domains.

In summary, we revealed distinct neural dynamics for shifting within vs. between perception and working-memory contents. We demonstrated how these distinct neural processes emerge quickly, evolve dynamically, are broadly distributed, and occur without delaying the timing of spatial orienting of attention. Besides successfully decoding the shift type, we were also able to decode the attentional domain both before and after the shift, thereby capturing the origin and destination of attention. Our results uncovered a direct link between the previously attended domain and the performed shift type, suggesting that lingering activity related to the first-cued domain affects processing following the second cue in a way that distinguishes within- from between-domain shifts. Brain imaging methods with higher spatial resolution should build on these findings to identify the brain networks that orchestrate shifts of attention between domains and investigate how they interact with the more established control networks involved in modulating shifts of attention within the external and internal domains.

## Methods and Materials

### Participants

The study was approved by the Central University Research Ethics Committee of the University of Oxford. The sample size was set to *n* = 25, based on a previous study from our lab targeting a similar research question via behavioural measures (17). To yield the targeted number of participants, we collected data from 26 participants. One participant was excluded following our a-priori behavioural trial-removal procedure (see Behavioural Analysis). Of the remaining 25 participants (age range: 19 to 36; mean age: 26.24; 14 female, 11 male), 22 reported being right-handed (three left-handed). Individuals provided written informed consent before participating in the study and were paid £15 per hour.

### Task and Procedure

The current task design builds on our previous behavioural study (17). In each trial, four randomly oriented bars appeared within coloured circular placeholders at quadrant locations. Early in the trial, two oriented bars were bilaterally presented and had to be encoded into working memory. Later in the trial, two bars appeared briefly in the remaining placeholders followed by masks. Between the initial and final presentation of the bars, one or two centrally presented colour cues signalled which of the four bars participants had to report by accurately reproducing its exact orientation. Cues that indicated a previously encoded bar directed attention internally (retro-cues or internal cues), indicating which item from working memory should be reported. In contrast, cues that indicated a previously unoccupied placeholder directed attention externally (pre-cues or external cues), specifying which item from the upcoming perceptual array should be reported.

At the start of each trial, a white central fixation cross (RGB value: [255, 255, 255]) surrounded by four coloured circular placeholders appeared against a black background (RGB value: [0, 0, 0]) for 750 ms. Each of the placeholders appeared in one of four highly distinguishable colours (RGB values: light blue [0, 159, 183], orange [217, 120, 0], green [47, 167, 0], pink [247, 35, 255]). The location mapping of the four placeholder colours was randomly determined on each trial. Placeholders subtended 3.5° visual angle and were centred in the top-left, top-right, bottom-left, and bottom-right at a distance of 4.5° visual angle from fixation. Placeholders stayed on the screen until the response was required at the end of the trial.

Next, two tilted grey bars (RGB value: [50, 50, 50]) subtending approximately 0.5°×3.4° appeared for 500 ms inside two of these placeholders (referred to as internal display). The orientation of the bars was randomly determined between 0 and 180°. Participants were instructed to memorise the orientation of these two bars. There were four possible configurations of the internal display, which occurred equally often throughout the experiment: the bars could appear in the placeholders located in the (1) top-left and top-right, (2) bottom-left and bottom-right, (3) top-left and bottom-right, and (4) top-right and bottom-left. As such, at the time of the internal display, there was always one bar presented in each visual hemifield.

The internal display was followed by a delay of 750 ms, during which only the fixation cross and placeholders remained on the screen. The central fixation cross then changed colour to match that of one of the placeholders for 250 ms. The colour change acted as an informative cue, indicating the potential item to be reported. This cue could indicate a placeholder that previously contained an item during the internal display (i.e., retro-cue) or a placeholder in which no item had been presented yet (i.e., an item of the upcoming external display; pre-cue). The cued location and attentional domain were pseudo-randomised such that each of the four placeholder locations and each domain was equally often indicated by the cue.

In 50% of the trials, the central fixation cross changed colour again after 1250 ms to match the colour of another placeholder for 250 ms (i.e., double-cue or shift trials). In the remaining 50% of the trials, only a single cue (i.e., single-cue or stay trials) was presented. In single-cue trials, the cue appeared at the same time as the first cue in double-cue trials and was 100% instructive, always indicating the target for report. Single-cue trials were included to increase the relevance of the first cue and ensure that participants shifted attention to the first-cued item rather than simply ignoring the first cue entirely. Single-cue trials served no other purpose for the behavioural and neural comparisons relevant to the research question addressed in the present study.

The double-cue trials were the important trials for our research question since these trials required a shift of attention. In particular, the double-cue condition was further divided into two sub-conditions: within- and between-domain shifts. In within-domain shift trials (i.e., external-to-external or internal-to-internal), the second cue indicated the alternative item within the same domain as the first cue occurring in the other hemifield. In between-domain shift trials (i.e., external-to-internal or internal-to-external), the second cue indicated the item in the alternative domain occurring in the other hemifield. Thus, in within- and between-domain shift trials, attention always needed to be reoriented from one hemifield to the other. The second-cued location and domain were pseudo-randomised such that each of the four placeholder locations and each domain were equally often indicated. In double-cue trials, the second cue instructed the target item for report with a certainty of 100%, thus overruling the first cue. Accordingly, the target item was always identical to the second-cued item. The colour of the first and second cue always differed in double-cue trials.

The single cue in single-cue trials and the second cue in double-cue trials were followed by an inter-stimulus interval of 1250 ms before the onset of a perceptual display (referred to as external display), containing two randomly tilted bars. The new items appeared in the yet-unoccupied placeholders for 50 ms. For example, if the bars in the internal display appeared in the top-left and bottom-right placeholders, bars in the external display occurred the top-right and bottom-left placeholders. To make the perceptual discrimination challenging, items in the external array were masked. The masking array consisted of four overlayed titled bars presented in each of the placeholders for 100 ms. At each mask location the four tilted bars differed 45° from the neighbouring bar. The overall orientation of each mask was randomly drawn at 0 ms and 50 ms, thus creating the impression of a dynamic display. The masking parameters were set to equate the difficulty of external and internal reports, based on piloting efforts.

The offset of the dynamic mask was followed by a 100-ms delay, after which a visual response dial was displayed, always starting in a random position. The response dial surrounding the fixation cross was the same size as the placeholders and included markers along a circle that corresponded to the ends of a bar. The dial rotated leftwards when pressing the left button and rightwards when pressing the right button of the response device held in the dominant hand. The dial rotated with a speed 1/10° per millisecond. Participants had unlimited time to complete the orientation report. When the dial aligned with the orientation of the target item, participants pressed either of the two buttons of the response device held in the non-dominant hand to confirm their response and continue with the task.

After a 100-ms delay in which only the fixation cross stayed on the screen, participants received feedback in the form of a number ranging from 0 to 100, with 100 indicating a perfect report and 0 indicating that the adjusted orientation was perpendicular to the angle of the target item. Feedback was presented 0.7° above the central fixation cross for 300 ms. Trials were separated by an inter-trial interval randomly drawn between 750 and 1000 ms. Between blocks, participants were presented with their average reproduction accuracy in the previous block.

The experiment consisted of 512 trials divided across 8 blocks, each including 64 trials. Of the total trials, 50% (256) were single-cue trials and 50% (256) were double-cue trials. Participants were equally likely to report an external or internal item in single-cue trials (128 each). Double-cue trials varied in the type of shift performed, that is, 50% (128) required a within-domain shift and the other 50% (128) a between-domain shift. Within-domain and between-domain shift trials were further split by the target domain. Ergo, the within-domain shift trials included 50% (64) external-to-external and 50% (64) internal-to-internal shifts, while the between-domain shift trials included 50% (64) external-to-internal and 50% (64) internal-to-external shifts. In each of the four unique shift conditions, participants were equally likely to shift their spatial attention between any of the eight possible cross-hemifield combinations of placeholders. The orthogonal manipulation of the to-be-reported target domain (external vs. internal) and the shift type (within-domain shift vs. between-domain shift) enabled us to independently investigate these two factors. Each block consisted of 50% (32) single- and 50% (32) double-cue trials.

To become familiarised with the procedure of the experiment, participants performed a practice block before starting the data collection. The whole experiment lasted approximately 65 min.

### Data Acquisition

We obtained whole-head MEG recordings in a magnetically shielded room using a 306-sensor MEG system (204 first-order planar gradiometers, 102 magnetometers; TRIUX neo, MEGIN OY, Espoo, Finland) at the Oxford Centre for Human Brain Activity. The MEG signal was sampled at 1000 Hz, with a high-pass filter at 0.1 Hz and a low-pass anti-aliasing filter at 330 Hz. Each experimental block was recorded separately, resulting in eight runs for most participants. However, due to technical issues, two individuals had fewer recordings, with six and seven recordings obtained, respectively.

Prior to data acquisition, we digitised the individual head shape with an electromagnetic position and orientation monitoring system (FASTRAK, Polhemus, Colchester Vermont, USA). Shapes included fiducial landmarks (nasion, right and left preauricular points) and about 250 additional points evenly spread on the participants’ scalp. Five Head Position Indicator (HPI) coils were placed on participants’ mastoid bones and forehead to keep track of participants’ head position inside the dewar through electromagnetic induction before and after each recording block.

Participants performed the task in a seated position. Participants were instructed to avoid head, body, and limb movements and to keep strict eye fixation during the experimental blocks. In addition to electrocardiography (ECG) and horizontal and vertical electrooculography (EOG), eye positions were continuously monitored by an MEG-compatible eye-tracking device with a sampling frequency of 1000 Hz (Eyelink 1000, SR-Research Ltd., Ottawa, Ontario, Canada). At the beginning of each experimental session, participants performed an eye-tracking calibration task to verify their gaze position on the screen. Calibration was repeated if drift was noticed during the experimental session.

We used the PsychoPy package (version 2021.2.3) (62) in Python for stimulus generation and stimulus delivery. The stimuli were projected on a translucent whiteboard using a DLP LED projector (ProPixx, VPixx Technologies Inc., Saint-Bruno, Quebec, Canada) at a 120-Hz refresh rate. The whiteboard was located at 120-cm distance from the participant, and it provided a projection area of 55×31-cm and 1920×1080-pixel resolution. A bimanual fibre-optic response device was used to collect manual responses.

Structural magnetic resonance imaging (MRI) scans were acquired for 22 participants at the Oxford Centre for Human Brain Activity using a Siemens Prisma 3T scanner, either immediately following the MEG recording or on a separate day. Existing structural 3T MRI scans for three participants were obtained from previous studies following data-sharing procedures.

### Behavioural Analysis

Behavioural data were analysed in R Studio (63). During pre-processing (similar to (17)) trials were removed when RTs (calculated from dial onset to response initiation) exceeded 5000 ms. Next, we removed trials for which the remaining RTs were 2.5 *SD* above the individual mean across all conditions. Moreover, we removed datasets with average reproduction errors (calculated by averaging the absolute difference between the angle of the target item and the reported angle) equal to or higher than 45° in any of the conditions (*n* = 1). After these exclusion steps, datasets from 25 participants remained in the main behavioural analysis, with an average of 16.56 ± 3.99 (*M* ± *SD*) trials removed. Additionally, we removed the bad MEG trials from the behavioural data analysis.

To meet parametric-test assumptions, the distribution and power coefficient of all dependent variables were inspected using the MASS package (64) and the Box-Cox procedure (65). As a result, RTs and reproduction errors were square-root-transformed. For the visualisation of RTs and reproduction errors, we used milliseconds and degrees as scales, respectively, while all behavioural inferential statistics were conducted on the square-root-transformed data.

The conditions of interest were target domain (having to report an external vs. internal item) and shift type (within vs. between). When comparing more than two means, we applied a repeated-measures analysis of variance (ANOVA) and reported η^2^_G_ as a measure of effect size. When evaluating only two means, we applied a paired samples *t*-test and report Cohen’s *d* as a measure of effect size.

### MEG Pre-processing

MEG data were analysed in Python using MNE-Python (version 1.4.2) (66) and OSL (version 1.1.0) (67) combined with custom code. For each individual run, we applied the Elekta MaxFilter (version 2.2) implementation of Spatiotemporal Signal Source Separation to remove external noise (TSSS) (68) to continuously compensate for head movements occurring during the run and to realign all runs per participant to a participant-specific head position. The reference head position for each participant was determined by minimising the head position distance between runs. Next, data were downsampled to 250 Hz and low-pass filtered at 50 Hz. Importantly, no high-pass filter was applied, since high-pass filtering can temporally displace multivariate information (69). Bad sensors were identified using a generalised extreme studentised deviate (ESD) test (70) at a 0.05 significance threshold and subsequently interpolated. On average, 1.62 ± 3.11 (*M* ± *SD*) bad gradiometers and 1.42 ± 2.02 bad magnetometers were identified and interpolated per run. We applied an independent component analysis (ICA) to regress out eye-movement and cardiac-related activity. Note that the maximum number of ICA components to be extracted was determined by the residual degrees of freedom after TSSS rank reduction during Maxfiltering (71). Data were epoched from -500 to 3000 ms relative to the first-cue onset without applying baseline correction (69). Next, we combined all individual runs for each participant by concatenating the data. Additionally, we remapped magnetometers onto gradiometers, allowing us to conduct all subsequent sensor-level analyses using a unified sensor type. Lastly, we employed an ESD trial rejection approach to detect and exclude trials with high variance. Per participant, 5.24 ± 6.96 (*M* ± *SD*) trials were removed. In addition, we removed the trials that were excluded in the behavioural data analysis. All following analyses were conducted using sensor-level MEG data, unless stated otherwise.

### Time-resolved Multivariate Pattern Analysis

We performed time-resolved multivariate analysis on broadband MEG data using the Scikit-learn toolbox (version 1.3.0) (72) and built-in decoding functions provided by MNE-Python (version 1.4.2) (66). The decoding pipeline closely followed the approach outlined in previous literature (73).

We extracted patterns of brain activity separately for each participant on a timepoint-by-timepoint basis (*mne.decoding.SlidingEstimator*). First, features (i.e., data at each time point for each sensor) were centred and scaled based on data from all trials (*sklearn.preprocessing.StandardScaler*). The true spatial dimensionality of the MEG data after the Maxfilter preprocessing algorithm is around 70, rather than the 204 gradiometer sensors (71). Therefore, to reduce the number of redundant features for decoding, we performed Principal Component Analysis on the standardised data while maintaining 99% of the variance (*sklearn.decomposition.PCA*). Training and testing were done on the same data using a 10-fold stratified cross-validation procedure (*sklearn.model_selection.StratifiedKFold*). First, trials were randomised and divided into 10 equal-sized folds. Next, a leave-one-out procedure was used on the 10 folds, such that the classifier was trained on 9 folds and tested on the remaining fold. This procedure was repeated 10 times until each fold was used once for testing. A LDA classifier (*sklearn.discriminant_analysis.LinearDisciminantAnalysis*) was utilised to train on the provided training data and labels. LDA is a supervised dimensionality-reduction technique that aims to find a linear combination of features that maximises class separability which has been recommended for neuroscientific decoding studies (34). We used the ROC-AUC as the scoring metric. This scoring metric takes into account the trade-off between true and false positive rates is considered a sensitive, nonparametric, and criterion-free measure of classification (31, 74). A ROC-AUC value of 0.5 means chance-level classification performance. The obtained classifier scores were averaged over all 10 folds, yielding a single decoding estimate per participant, timepoint, and condition. Lastly, decoding time courses were smoothed with a one-dimensional Gaussian filter with a *SD* of 10 samples (i.e., 40 ms).

In addition, we used temporal-generalisation methods (35) to examine whether there are recurrent neural activation patterns over time, that is, whether an activation pattern observed at one timepoint is observed again at a later timepoint. Generalisation off the diagonal in the temporal generalisation matrix is driven by overlap in the activation pattern at two different timepoints, and therefore implies that the activation pattern recurred. Temporal generalisation allows us to examine such potential overlaps in representations over time. This approach results in timepoint-by-timepoint decoding matrices in which each cell corresponds to a classification accuracy at a unique training- and test-timepoint combination.

To test which brain areas contributed most to the classifier likelihoods observed in our multivariate methods, we ran a sensor-space searchlight decoding analysis (similar to 13, 30, 34, 73), analogous to the searchlight analysis developed for fMRI (76). For this analysis, we ran the same analysis as outlined above across small clusters of 15 neighbouring MEG sensors, resulting in decoding scores for each sensor and timepoint. These decoding scores were averaged into time clusters of 300 ms each, resulting in 10 topographical maps between 0 and 3000 ms. These topographical plots show whether decoding results were primarily driven by specific sensor clusters. Note that although this approach differs from the analysis of activation patterns, it ultimately serves the same purpose: identifying the sources of neural processes in space and time (77). We did not test any specific hypotheses regarding the spatial distributions of the effects and therefore did not run any statistical tests on the results

Even though stratified cross-validation is highly sensitive to class imbalances (31), we additionally applied between-class balancing using under-sampling prior to the decoding analysis, such that the number of trials was equal across the classes. This further ensured that during training the classifier would not only achieve a high decoding performance merely by systematically and blindly voting for the majority class (78). Note that as the design was balanced in terms of trial counts, the between-class balancing was employed to account for small imbalances due to trial-rejection procedures used for data cleaning.

### Relationship Between Decoding and Behaviour

To investigate the relationship between the observed decoding of within- vs. between-domain shifts of attention and the behavioural shift costs, we employed the following approach. Firstly, we identified trials with low and high classifier performance for each participant by computing timepoint-by-timepoint decoding scores and averaging them across the interval of interest. Subsequently, we performed a median split based on these average decoding scores, creating two groups of trials: one with low decoding scores and the other with high decoding scores. We then included the median-split factor in our behavioural analysis when comparing within- to between-domain shifts.

### Control Analysis: Eye Movements

Involuntary eye movements are a common problematic covariate in studies using multivariate approaches (32, 33). To test for an influence of gaze on the MEG findings, we ran the same MEG decoding analyses on our eye-tracking data. We acquired a total of 23 eye-tracking datasets, consisting of 15 binocular recordings and 8 monocular recordings. For the binocular recordings, we merged the x- and y-coordinates from both eyes to create a single metric. The data were then epoched from -500 ms to 3000 ms relative to the first-cue onset and were baseline-corrected prior to cue onset (-500 to 0 ms). Next, we trained the classifier to distinguish between different conditions at each timepoint using only the x- and y-coordinates of the eye-tracking data. Instead of a LDA classifier, we used a Support Vector Machine classifier (SVC; *sklearn.svm.SVC*) with Radial Basis Function (RBF) kernel. The reason behind opting for SVC with RBF kernel is the likelihood that the distribution of x- and y-coordinates of gaze is not linearly separable. In all conditions, participants were instructed to maintain fixation, making it challenging to distinguish between different classes or conditions using a simple linear boundary. The RBF kernel is a powerful tool for transforming non-linearly separable data into a higher-dimensional space, making it possible to separate the data using a hyperplane (79). This method is particularly effective when dealing with datasets a smaller number of features than of samples.

By computing mutual information scores over correctly classified MEG trials on a timepoint-by-timepoint basis, we assessed how much information was shared between the MEG decoding results and eye-movement decoding results (33). Mutual information quantifies the extent to which knowledge of one variable reduces uncertainty about another variable. Higher mutual information values indicate a stronger dependence between two variables, suggesting that changes in one variable are informative about the other.

### Time-frequency Analysis

We generated time-frequency representations of power by convolving the data with Morlet wavelets across the frequency range of 3 to 40 Hz. For each frequency, we used a fixed 300-ms time window such that the number of cycles changed with the frequency. The power time series in the planar gradiometer pairs were then combined (i.e., root mean square), resulting in a 102-sensor combined planar gradiometer map in sensor space.

For the comparison between contra- and ipsilateral power, we computed activity following left- and right-directing cues, separately for posterior left sensor clusters (MEG1632+1633, MEG1642+1643, MEG1912+1913, MEG1922+1923, MEG1942+1943, MEG2042+2043) and right sensor clusters (MEG2032+2033, MEG2312+2313, MEG2322+2323, MEG2342+2343, MEG2432+2433, MEG2442+2443), and subsequently pooled the contra- vs. ipsilateral contrasts between them. We expressed the contra-vs-ipsi contrast as a normalised difference [i.e., ((contra – ipsi)/(contra + ipsi)) × 100]. Sensors were chosen based on prior MEG studies from our lab that have consistently implicated the same set of posterior sensors for capturing neural activity related to lateralised visual stimuli (80, 81). While we selected these sensor clusters based on independent data, we could confirm their validity and appropriateness in the current data (Fig. 5g and h overlay of these sensor clusters on the relevant left vs. right topographies).

To extract time courses of alpha lateralisation, we averaged the contra- vs. ipsilateral activity across the predefined alpha band (8-12 Hz) in the visual sensors. Topographies of the lateralised visual activity were obtained by contrasting trials in which cues indicated visual content in the right vs. left hemifield. To depict the topographies associated with relevant condition comparisons, we focused on the alpha frequency bands between 0 and 3000 ms in predefined steps of 500 ms.

### Event-related-fields Analysis

To complement the decoding and time-frequency analyses, we performed a univariate analysis of ERFs. The sensor-level MEG data used in the ERF analysis underwent similar preprocessing as the data for the remaining analyses, with two important differences. First, to eliminate slow drifts, we applied a 1-Hz high-pass filter to the continuous data. Second, epochs were baseline-corrected using the 500-ms window preceding the onset of the first cue. As with the time-frequency analysis, planar gradiometer pairs were combined using the root mean square, resulting in a 102-sensor combined planar gradiometer map in sensor space.

The combined planar gradiometers were divided by hemisphere (left and right) and lobe (frontal, temporal, parietal, and occipital) into eight sensor clusters with 12 or 13 pairs each (Appendix SI, Table S1) according to (82). For each sensor cluster and shift-type condition, we computed the averaged ERF time course.

Moreover, to compare lateralised visual activity in posterior regions, we calculated separate waveforms from the left and right sensor clusters (using the same sensor clusters as in the time-frequency analysis), following both left- and right-directing cues. This resulted in four separate waveforms: two from the left sensors (i.e., cue-left, cue-right) and two from the right sensors (i.e., cue-left, cue-right]). Contra- and ipsilateral ERF time courses were then calculated by averaging the waveforms of trials wherein the cue indicated an item appearing on the opposing or same side as the relative sensor cluster, respectively. To compare lateralised brain activity between shift types, we computed the contra- vs. ipsilateral contrast for each condition. All ERF time courses were smoothed with a one-dimensional Gaussian filter with a *SD* of 5 samples (i.e., 20 ms).

### Source Reconstruction

To visualise sources of oscillatory brain activity, sensor-level MEG scans were source reconstructed using OSL (version 1.1.0) (67). For each participant, structural MRI and MEG coordinate systems were co-registered by matching digitised anatomical fiducial landmarks and head shape points on the participant-specific T1 scans. Forward modelling used a single layer representing the inner skull surface (i.e., single-shell boundary element model). The epoched sensor-level data were reconstructed onto a 5-mm isotropic grid using a unit-noise-gain Linearly Constrained Minimum Variance (LCMV) beamformer (71, 83). The data covariance matrix used to calculate the beamformer was estimated using the continuous pre-processed sensor-level data for each participant, regularised to a rank of 60 using PCA. A diagonal matrix containing the variance of each sensor type was used as noise covariance. The beamformer provides the activity in the x-, y-, z-direction at each grid point. This was projected onto an axis that maximises power to obtain a single time course for each grid point (i.e., voxel).

The participant-averaged voxelwise difference in alpha-band activity between trials in which cues indicated visual content in the right vs. left hemifield was plotted on the MNI152 standard brain template.

### Statistical Analysis

Statistical evaluation of MEG data was conducted using a cluster-based non-parametric permutation approach with 1024 permutations and a cluster alpha of 0.05. This approach offers a solution to the challenges related to multiple comparisons in the statistical analysis of MEG data, especially when dealing with a large number of comparisons. In particular, this approach evaluates temporal clusters observed in the original (non-permuted) data against a permutation distribution derived from the largest temporal cluster found after each permutation of the condition labels (84).

In a supplementary analysis, we additionally utilised a jack-knife approach to identify differences in decoding-onset times. For the jack-knife approach, we computed decoding time-course averages by excluding each of the *n* data sets once, allowing us to obtain *n* leave-one-out grand averages. Next, we identified the time point at which the decoding time course reached 50% of its peak. Jack-knifed standard errors, *t-*values, and *p*-values were adjusted according to (85).

## Author Contributions

Daniela Gresch: Conceptualization, Methodology, Software, Investigation, Formal analysis, Visualization, Writing – original draft. Sage E.P. Boettcher: Conceptualization, Methodology, Writing – review & editing, Funding acquisition, Supervision. Chetan Gohil: Formal analysis. Freek van Ede: Conceptualization, Methodology, Writing – review & editing, Funding acquisition, Supervision. Anna C. Nobre: Conceptualization, Methodology, Writing – review & editing, Funding acquisition, Supervision.

## Acknowledgements

This research was funded by a Wellcome Trust Senior Investigator Award (104571/Z/14/Z) and a James S. McDonnell Foundation Understanding Human Cognition Collaborative Award (220020448) awarded to A.C.N.; an Experimental Psychology Society Postdoctoral Fellowship awarded to S.E.P.B.; and an ERC Starting Grant from the European Research Council (MEMTICIPATION, 850636) and NWO Vidi Grant from the Dutch Research Council (14721) awarded to F.v.E; and a Wellcome Trust grant supporting C.G. (215573/Z/19/Z). The research was also supported by the NIHR Oxford Health Biomedical Research Centre. The Wellcome Centre for Integrative Neuroimaging is supported by core funding from the Wellcome Trust (203139/Z/16/Z). For the purpose of open access, the authors have applied a CC-BY public copyright licence to any Author Accepted Manuscript version arising from this submission.

## Competing Interests

The authors declare no competing interests.

## Supporting Information

**Fig. S2.**
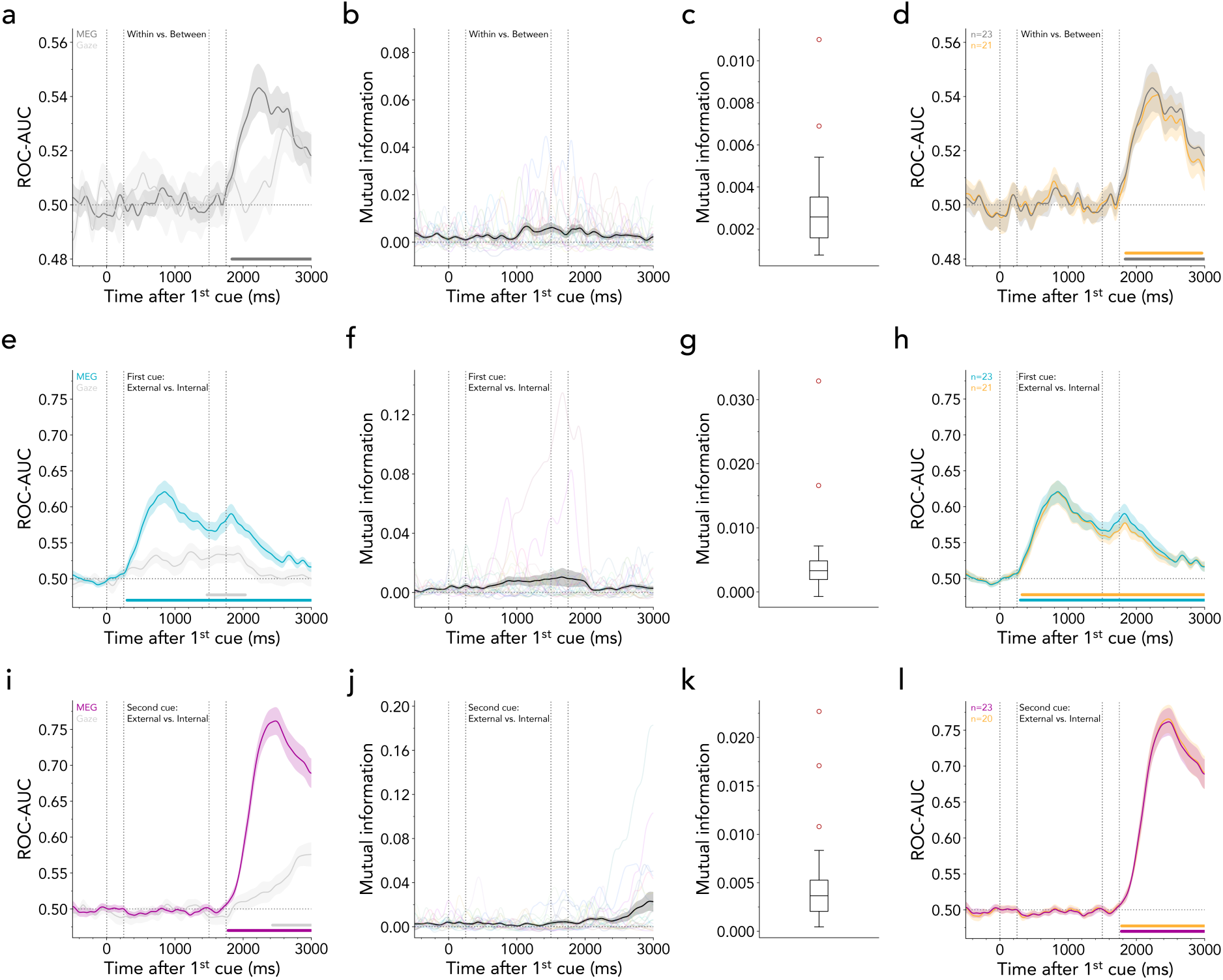
Eye movements do not account for MEG decoding results. **(a)** Average time-resolved decoding of within- vs. between-domain shifts based on MEG and eye-tracking data. Only MEG data sets were included that had an accompanying eye-tracking data set (*n* = 23). Horizontal lines indicate significant clusters (MEG decoding: ∼1840-3000 ms, cluster *p* < 0.001). Time course shows *M* ± *SEM* across participants. **(b)** Average mutual information between MEG and eye-tracking data for the within- vs. between-shift decoding. Faded lines indicate individual participants. **(c)** Box-and-whisker plot according to (1) displaying distribution of average mutual information across -500 to 3000 ms relative to first-cue onset. Red dots represent outliers. **(d)** Average time-resolved MEG decoding of within- vs. between-domain shifts after removing outliers (*n=*23: ∼1840-3000 ms, cluster *p* < 0.001, *n=*21: ∼1840-3000 ms, cluster *p* < 0.001). **(e)** Same as (a) but for external vs. internal first cues (MEG decoding: ∼300-3000 ms, cluster *p* < 0.001, eye decoding: ∼1470-2030 ms, cluster *p* = 0.042). **(f)** Same as (b) but for external vs. internal first cues. **(g)** Same as (c) but for external vs. internal first cues. **(h)** Same as (d) but for external vs. internal first cues (*n=*23: ∼300-3000 ms, cluster *p* < 0.001, *n=*21: ∼320-3000 ms, cluster *p* < 0.001). **(i)** Same as (a) but for external vs. internal second cues (MEG decoding: ∼1780-3000 ms, cluster *p* < 0.001, eye decoding: ∼2430-3000 ms, cluster *p* = 0.010). **(j)** Same as (b) but for external vs. internal second cues. **(k)** Same as (c) but for external vs. internal second cues. **(l)** Same as (d) but for external vs. internal second cues (*n=*23: ∼1780-3000 ms, cluster *p* < 0.001, *n=*20: ∼1780-3000 ms, cluster *p* < 0.001).

**Fig. S3.**
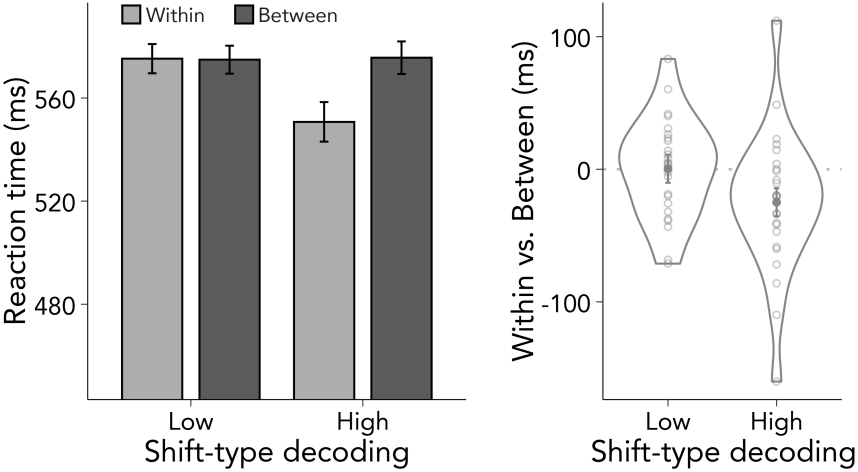
Within- vs. between-shift decoding is behaviourally relevant. Left panel: Reaction times (RTs) as a function of shift type (within vs. between) and median split (low vs. high). Error bars represent *SEM*. Right panel: Shift cost in RTs defined as the difference of within- and between-shift trials for each half of decoding strength. Dots represent individual participants.

**Fig. S4.**
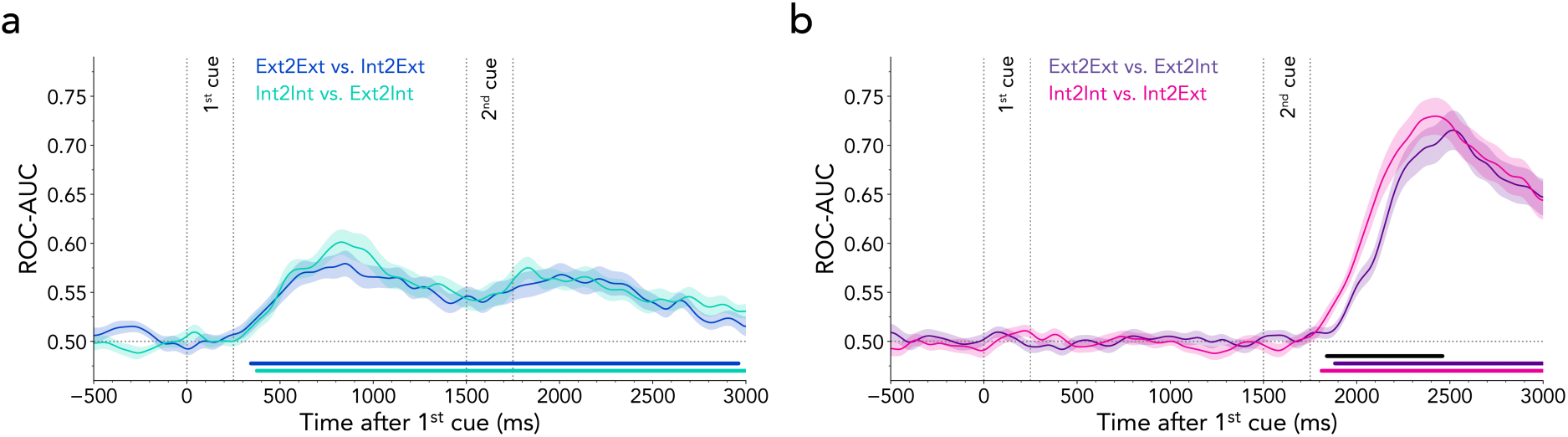
Time-resolved MEG decoding of different trial types. **(a)** Average time-resolved classifier performance for distinguishing external-to-external from internal-to-external shifts, and internal-to-internal from external-to-internal shifts. Time courses show *M* ± *SEM* across participants. Coloured horizontal lines indicate significant clusters (external-to-external vs. internal-to-external cluster: ∼340-2960 ms, cluster *p* < 0.001; internal-to-internal vs. internal-to-external cluster: ∼380-3000 ms, cluster *p* < 0.001). **(b)** Same as (a) but decoding internal-to-internal vs. internal-to-external shifts, and external-to-external vs. external-to-internal shifts (external-to-external vs. external-to-internal cluster: ∼1880-3000 ms, cluster *p* < 0.001; internal-to-internal vs. internal-to-external cluster: ∼1810-3000 ms, cluster *p* < 0.001). Black horizontal line denotes significant difference between decoding time courses (∼1840-2460 ms, cluster *p* < 0.001). Since statements regarding the exact onset of an effect are not supported by permutation tests (2), we quantified the timings of the decoding time courses more formally by applying a jack-knife approach to compare the point at which the decoding time courses reached 50% of their peak value. This revealed significantly earlier onset times for internal-to-internal vs. internal-to-external decoding (*M* = 2046.240, *SEM* = 20.800) as compared to external-to-external vs. external-to-internal decoding (*M* = 2133.92, *SEM* = 16.738; *t*_(24)_ = -3.379, *p* = 0.002).

**Fig. S5.**
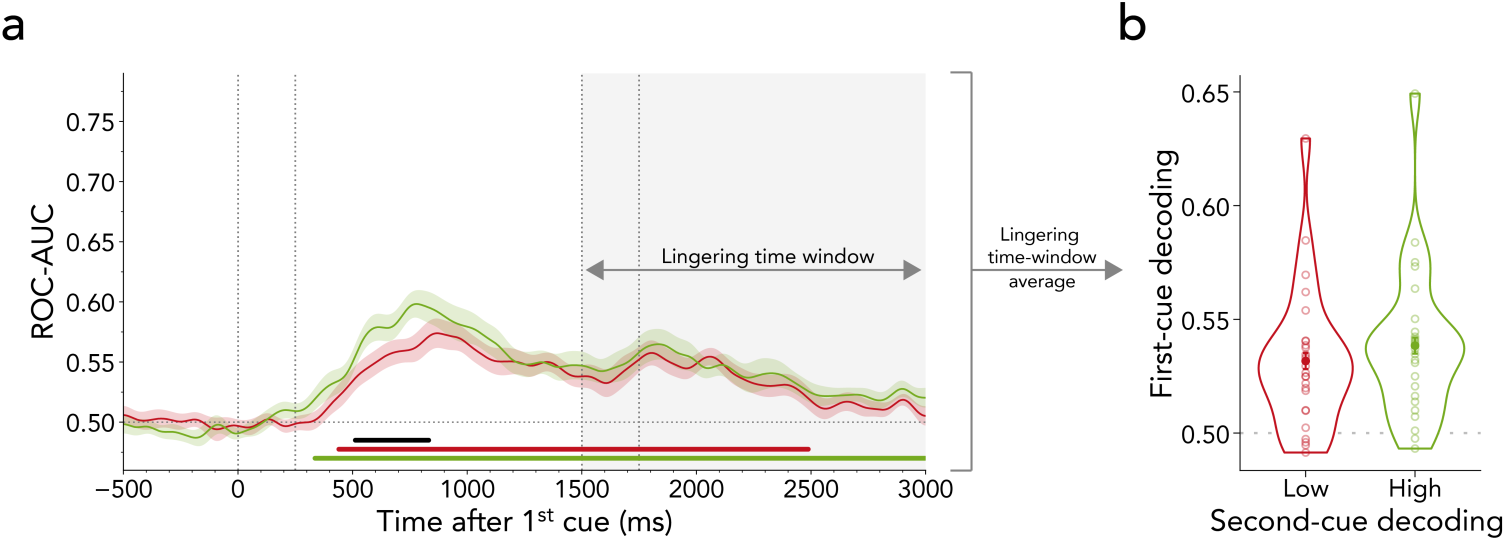
No evidence for a trade-off in decodability between first- and second-cued attentional domains. **(a)** We median-split the MEG data into trials with low vs. high decoding performance in distinguishing the second-cued domain following the second cue (i.e., 0-1500 ms after second-cue onset). Next, we trained a classifier separately on both data sets to predict the first-cued domain (external vs. internal). In trials with low and high second-cue decoding, we could successfully predict the first-cued attentional domain after first-cue appearance and the decodability remained significant following the second cue (low decoding: ∼440-2490 ms, cluster *p* < 0.001; high decoding: ∼340-3000 ms, cluster *p* < 0.001). We did not observe any trade-off in the decoding of the first-cued attentional state between low vs. high second-cue decoding following the second cue (i.e., grey area). There was an earlier difference in the decodability of external vs. internal first-cue trials when the decodability of the second cue was low vs. high (∼510-830 ms, cluster *p* = 0.012). Time course shows *M* ± *SEM* across participants. **(b)** Average first- cued domain decoding in the second-cued period (i.e. grey area in (a)) as a function of low vs. high second-cued domain decoding. There was no difference between low and high second-cue decoding in predicting the first-cued domain (*t*_(24)_ = 1.348, *p* = 0. 190, *d* = 0.270). Error bars represent *SEM*.

**Fig. S6.**
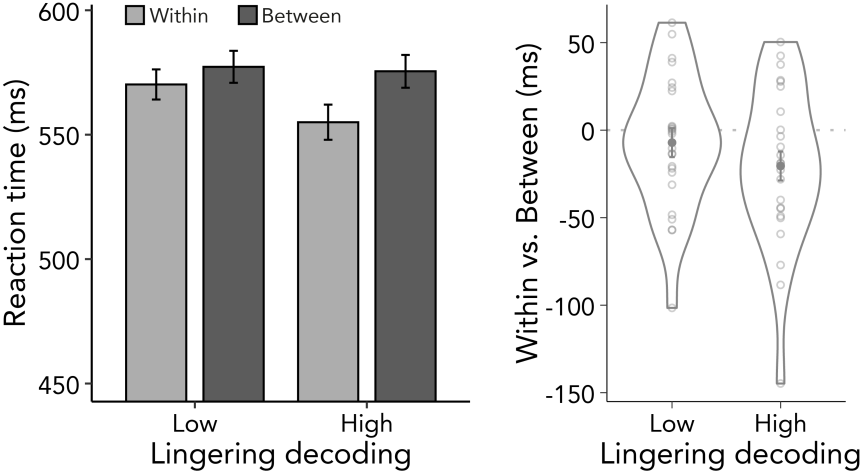
Lingering first-cue activity is behaviourally relevant. We median-split the MEG data into trials with low vs. high decoding performance in distinguishing the first-cued domain following the second cue (i.e., 0-1500 ms after second-cue onset). Next, we included the median-split factor in our behavioural analysis when comparing within- to between-domain shifts. Left panel: Reaction times (RTs) as a function of shift type (within vs. between) and median split (low vs. high). Error bars represent *SEM*. Right panel: Shift cost in RTs defined as the difference of within- and between-shift trials for each half of decoding strength. Dots represent individual participants.

**Fig. S7.**
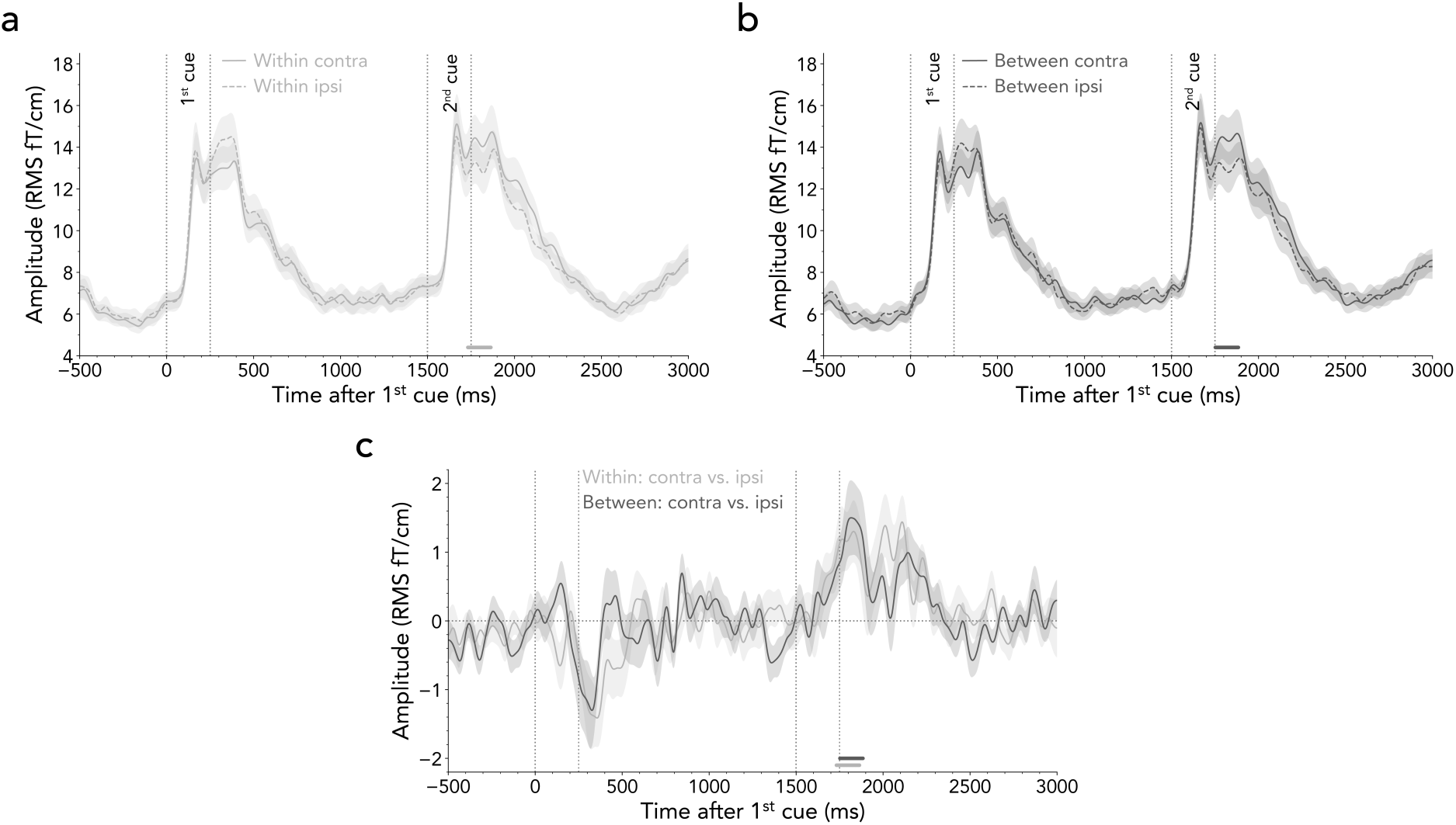
Lateralised visual activity in posterior regions. **(a)** Event-related fields (ERFs) for within-domain shift trials from combined posterior gradiometers contra- and ipsilateral to the second-cued hemifield. The dashed and solid line depict the contra- and ipsilateral ERF, respectively. Time course shows *M* ± *SEM* across participants. The difference between contra- vs. ipsilateral lines is the equivalent of the N2pc which is typically observed in electroencephalography studies employing visual-spatial attention tasks. Horizontal lines indicate significant clusters when comparing contra- and ipsilateral ERFs (∼1730-1870 ms, cluster *p* = 0.048). **(b)** Same as (a) but for between-domain shift trials (∼1750-1880 ms, cluster *p* = 0.027). **(c)** Contra- vs. ipsilateral contrast for within- and between-domain shifts. There was no significant difference between conditions.

**Table S1.** Sensor grouping for event-related fields. For univariate analyses of event-related fields, gradiometer pairs were categorised into eight groups according to different brain regions. Each of the channel groups consists of 12 or 13 sensor pairs.

|  | Left hemisphere | Right hemisphere |
| --- | --- | --- |
| **Frontal lobe** | MEG0622+0623, MEG0332+0333,<br>MEG0322+0323, MEG0342+0343,<br>MEG0122+0123, MEG0612+0613,<br>MEG0542+0543, MEG0312+0313,<br>MEG0822+0823, MEG0532+0533,<br>MEG0522+0523, MEG0512+0513,<br>MEG0642+0643 | MEG1412+1413, MEG1222+1223,<br>MEG1232+1233, MEG1242+1243,<br>MEG1032+1033, MEG1012+1013,<br>MEG1022+1023, MEG0932+0933,<br>MEG1212+1213, MEG0942+0943,<br>MEG0922+0923, MEG0912+0913,<br>MEG0812+0813 |
| **Temporal lobe** | MEG0222+0223, MEG0212+0213,<br>MEG0132+0133, MEG0232+0233,<br>MEG0242+0243, MEG1542+1543,<br>MEG1512+1513, MEG1622+1623,<br>MEG1612+1613, MEG1522+1523,<br>MEG1532+1533, MEG0112+0113,<br>MEG0142+0143 | MEG1312+1313, MEG1322+1323,<br>MEG1442+1443, MEG1432+1433,<br>MEG1342+1343, MEG1332+1333,<br>MEG2612+2613, MEG2412+2413,<br>MEG2422+2423, MEG2642+2643,<br>MEG2632+2633, MEG1422+1423,<br>MEG2622+2623 |
| **Parietal lobe** | MEG0712+0713, MEG0632+0633,<br>MEG0432+0433, MEG0422+0423,<br>MEG0442+0443, MEG0412+0413,<br>MEG0742+0743, MEG1832+1833,<br>MEG2012+2013, MEG1822+1823,<br>MEG1842+1843, MEG1812+1813,<br>MEG1632+1633 | MEG0732+0733, MEG2242+2243,<br>MEG2022+2023, MEG2212+2213,<br>MEG2232+2233, MEG2222+2223,<br>MEG2442+2443, MEG0722+0723,<br>MEG1042+1043, MEG1142+1143,<br>MEG1112+1113, MEG1122+1123,<br>MEG1132+1133 |
| **Occipital lobe** | MEG2042+2043, MEG1912+1913,<br>MEG1942+1943, MEG1642+1643,<br>MEG1722+1723, MEG1712+1713,<br>MEG1732+1733, MEG1922+1923,<br>MEG2112+2113, MEG1932+1933,<br>MEG1742+1743, MEG2142+2143 | MEG2032+2033, MEG2342+2343,<br>MEG2132+2133, MEG2332+2333,<br>MEG2542+2543, MEG2512+2513,<br>MEG2322+2323, MEG2312+2313,<br>MEG2532+2533, MEG2432+2433,<br>MEG2522+2523, MEG2122+2123 |

## Notes

### Competing Interest Statement

The authors have declared no competing interest.

### Summary of Updates

The manuscript has been revised based on the feedback from the reviewers.

